# Energy-Regularized Graph Learning for Multiscale Spatial Representation

**DOI:** 10.1101/2025.08.22.671846

**Authors:** Qiyu Gong, Kai Cao, Jackson A. Weir, Qitian Wu, Dongshunyi Li, Fei Chen

## Abstract

Spatial omics map gene and protein expression *in situ*, demanding methods that recover cellular and tissue architecture from noisy, high-dimensional data. This need spans two scales: (i) subcellular, where high-resolution measurements must be grouped into coherent cells, and (ii) tissue-wide, where the goal is to recover spatial domains. Existing embedding approaches either ignore space or rely on static neighborhood graphs that over- or under-smooth local heterogeneity. We present Glimmer (**G**raph-**L**earned **I**nference of **M**ultiscale **M**olecular **E**mbedding for Spatial **R**epresentation), a unified framework that learns adaptive neighborhood graphs by minimizing Dirichlet energy under a log-barrier regularizer. Beginning from a *k*-nearest neighbor scaffold, Glimmer adaptively reweights edges to balance molecular similarity and spatial proximity, yielding interpretable, locally smoothed embeddings. These graphs enable segmentation-free reconstruction from transcript localizations or fine bins and support the discovery of tissue niches at large tissue scales. Across diverse datasets and modalities (Slide-tags, Slide-seq, MERFISH, Xenium, and CODEX), Glimmer surpasses kernel and graph neural network based methods in clustering accuracy and spatial conservation. At the subcellular level, Glimmer corrects transcript-to-cluster misassignments on lymph node slides, thereby improving gene specificity within biological clusters and addressing a key challenge in spatial transcriptomics. At the tissue-wide scale, Glimmer enables accurate region identification, as demonstrated in tonsil tissue by resolving germinal center subregions, which in turn facilitates niche-specific immune profiling. Glimmer thus offers a generalizable framework for spatial representation learning across scales and modalities, enabling comprehensive insights into tissue architecture and cellular ecosystems.

## Introduction

The complexity and multifunctionality of tissues arise from the spatial organization of diverse cellular communities across biological scales^1^. Spatial omics technologies — including imaging-based methods (e.g., MERFISH^2^, CODEX^3^, Xenium^4^) and sequencing-based approaches (e.g., Slide-seq^5^, Slide-tags^6^, Visium) — have transformed tissue biology by mapping molecular profiles within native spatial contexts. However, these data are inherently high-dimensional, sparse, and noisy due to biological complexity and technical limitations, especially in spatial transcriptomics where dropout and redundancy are prevalent^7^. To extract meaningful biological patterns, it is critical to embed the data into denoised, low-dimensional spaces that support downstream clustering^8,9^.

Traditional methods such as Principal Component Analysis (PCA)^10^ and Non-negative Matrix Factorization (NMF)^11^ ignore spatial context, potentially missing spatial structures or misclassifying cell states shaped by local microenvironment. Recent approaches address this by integrating spatial and molecular information to better capture local structure^12^. However, balancing these two signals remains challenging: overemphasizing expression may obscure spatial patterns, whereas over-relying on spatial proximity can blur cellular heterogeneity. Navigating this trade-off is essential for capturing the tissue complexity across scales^13,14^.

Recent advances in spatial clustering have led to two major paradigms: kernel-based approaches (e.g., BANKSY^15^, SPIN^16^, SpatialPCA^17^, GraphPCA^18^) and graph neural network (GNN)-based models (e.g., STAGATE^19^, GraphST^20^, SpaceFlow^21^). Kernel-based methods integrate spatial and molecular features using fixed neighbor aggregation strategies. For instance, BANKSY employs azimuthal filtering to capture expression gradients; SPIN adopts random sampling neighbors; SpatialPCA encodes spatial proximity using a Gaussian kernel within a probabilistic PCA framework; and GraphPCA enforces spatial smoothness via a graph Laplacian penalty. These methods are efficient and scalable, but the fixed, globally shared kernels limit their flexibility in capturing the complex local heterogeneity experienced by individual cells. In contrast, GNN-based models integrate spatial and molecular features through learnable aggregation, allowing adaptive information flow across spatial neighborhoods. For example, STAGATE introduces attention mechanisms^22^ to adaptively weigh neighboring nodes, while GraphST and SpaceFlow further enhance spatial modeling through contrastive learning and graph refinement. Although GNNs offer flexible embeddings, they are susceptible to ill-posed objectives in label-scarce settings and are limited by hyperparameter sensitivity and scalability challenges^23–25^. This trade-off between scalability and flexibility motivates combining the adaptability of GNNs with the efficiency of kernel-based frameworks.

While existing methods primarily focus on niche-level clustering, recent advances in imaging-based spatial omics have extended the task to the subcellular resolution. Despite differences in scale, the common goal remains: embedding molecular profiles into spatially structured latent spaces. However, subcellular clustering poses unique challenges, as it demands accurate transcript assignment to preserve intracellular coherence and distinguish intercellular heterogeneity^26–28^. Segmentation-based methods often suffer from boundary inaccuracies and transcript-image mismatches, while segmentation-free approaches like FICTURE^29^ alleviate these issues but operate at lower resolution, which limits their ability to capture cellular details. These challenges underscore the need for adaptive embeddings that integrate molecular and spatial features across scales to support transcript-level clustering within cells.

Capturing the organizational complexity of tissues across scales requires adaptive embedding strategies^30^. While existing methods perform well at specific levels, they are often task-specific and lack explicit mechanisms to balance spatial proximity and molecular similarity across diverse biological contexts. For example, in structured organs like the brain, minimal spatial smoothing may suffice as expression aligns with anatomy^31^, whereas disordered systems such as tumors benefit from stronger spatial regularization to overcome noisy neighbor compositions^32^. At subcellular or single-cell resolution, embeddings must preserve fine-grained expression differences while maintaining spatial coherence^33^. However, fixed neighbors and static weights in current methods limit adaptability across tissues and tasks, while rigid losses and unclear aggregation reduce interpretability and generalization. These issues point to the need for a flexible, context-aware framework that learns to integrate spatial and molecular cues in a data-driven way.

To address these challenges, we introduce Glimmer, a unified representation learning framework that balances spatial proximity and molecular similarity by learning context-aware neighborhood graphs through Dirichlet energy minimization^34^. With built-in neighbor weight regularization and sparsity control, Glimmer constructs localized graphs for flexible and interpretable spatial embeddings. It generalizes across omics modalities, platforms, and resolutions, supporting both tissue-scale domain segmentation and subcellular transcript-level clustering. The localized graph design further facilitates robust cross-dataset integration, capturing both conserved and context-specific clusters. Applied to Slide-tags, Slide-seq, MERFISH, and Xenium datasets, Glimmer consistently outperforms state-of-the-art methods in clustering accuracy and spatial coherence. In human tonsil, it delineates germinal center zones and reveals niche-specific signaling, linking cellular states to tissue architecture. Its segmentation-free design further supports accurate transcript-to-cluster assignment at subcellular resolution. Together, these results establish Glimmer as a scalable, interpretable, and biologically grounded framework for multiscale spatial clustering.

## Result

### Glimmer introduces a learnable graph-based framework for spatial omics data representation

Achieving biologically meaningful representations in spatial omics remains challenging, as results are highly sensitive to how spatial neighboring cells are selected and weighted. Such representations can be used as a basis for the discovery of spatial gene expression variability and tissue niches. Learning useful representations relies on both molecular similarity and spatial proximity. When molecular similarity is allowed to dominate, the analysis tends to split tissue into salt-and-pepper clusters that are transcriptionally coherent yet spatially fragmented; when physical distance takes precedence, clusters become geometry-driven and mask functional diversity. A method that balances these opposing aspects is essential for revealing genuine tissue niches while preserving the single-cell resolution of modern assays (**Fig. 1a**).

**Figure 1.**
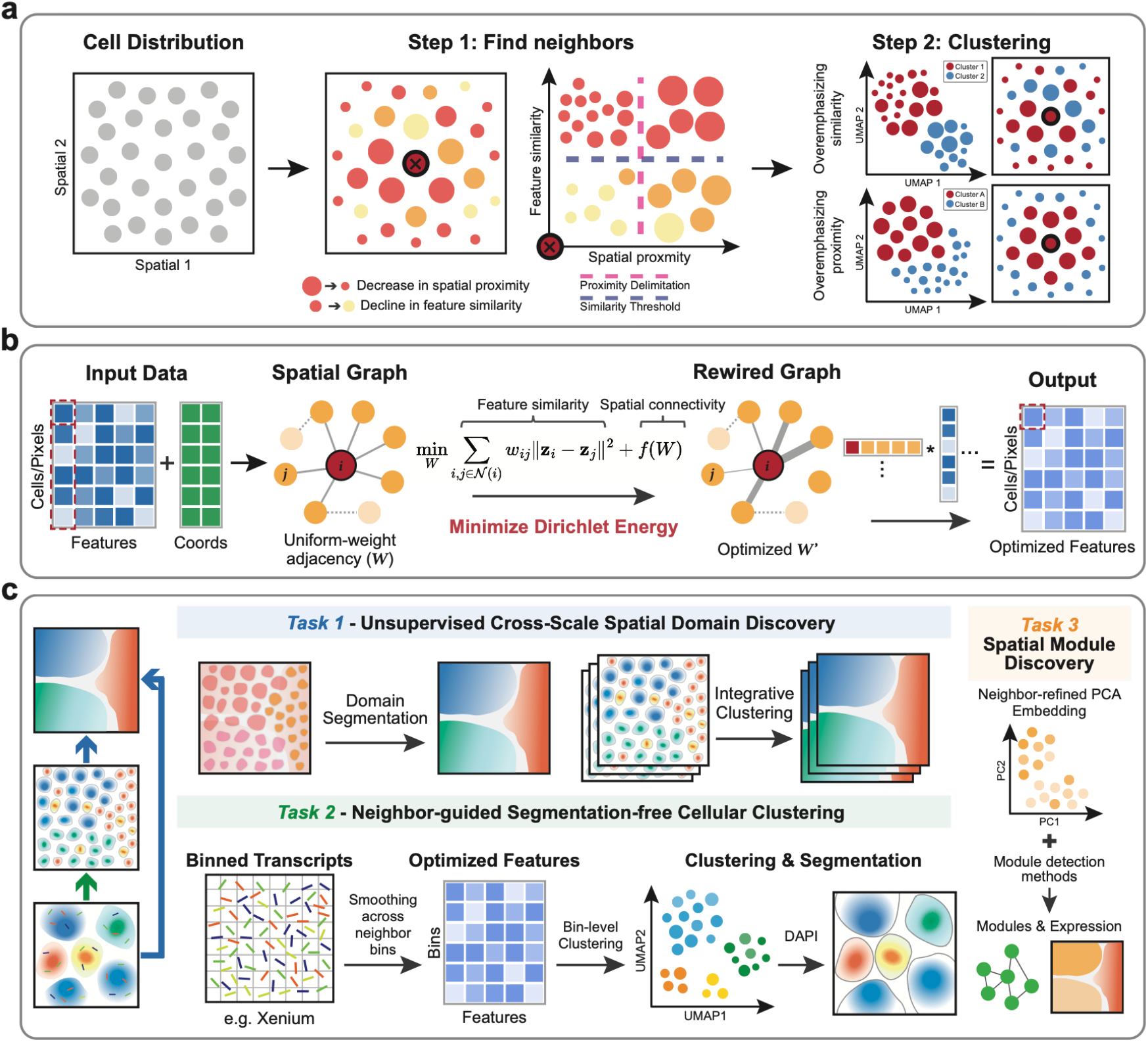
Overview of the Glimmer framework for multiscale spatial representation learning and downstream applications. **(a)** *Conceptual framework*. Glimmer refines spatially informed representations by constructing a neighbor graph that balances molecular similarity and spatial proximity. Left: Cells positioned in the physical space. Middle: Neighborhoods defined by dual thresholds on spatial and transcriptional similarity, preserving coherent local structure while filtering spatially close but functionally distinct neighbors. Right: Embedding informed by different neighbor criteria results in different clustering behavior—favoring molecular similarity can miss spatial coherence, while overemphasizing proximity may group dissimilar cells. **(b)** *Implementation*. Given input features and coordinates, a uniform spatial graph is first built. Through Dirichlet energy minimization, the graph is reweighted to balance transcriptionally and spatially coherent neighbors. The optimized graph supports weighted aggregation to generate refined, context-aware embeddings. **(c)** *Applications*. Task 1: Multi-scale spatial domain discovery within samples, with neighbor-refined embeddings supporting integrative clustering across slides. Task 2: Segmentation-free analysis of spatial transcriptomic bins (e.g., Xenium) via neighbor smoothing to derive cell-like features for unsupervised cellular clustering. Task 3: Spatial module discovery using neighbor-refined embeddings followed by spatial co-expression analysis.

Glimmer addresses this challenge by optimizing the neighbourhood graph by casting graph construction as a Dirichlet-energy minimization problem with two opposing terms (**Fig. 1b**). The first term, feature-similarity preservation, minimizes transcriptomic dissimilarity with spatially decayed weights, thereby reinforcing connections between spatially close and expression-similar cells. The second term, spatial-proximity regularisation, adds a logarithmic barrier that prevents edge weights from vanishing, thus promoting local smoothness. Starting from a uniform nearest-neighbour scaffold, Glimmer iteratively reweights edges via minimization of Dirichlet energy. Mathematically, Dirichlet energy quantifies the degree of variation in a function over a graph; minimizing it encourages smoothness across neighboring nodes^34,35^.

The learned graph yields locally smoothed embeddings in which spatial neighbours share similar molecular profiles while real biological gradients remain intact. At the tissue level, this enables the identification of coherent functional regions defined by recurrent patterns of molecular cooperation. At the cellular level, because Glimmer operates directly on fine spatial bins, it enables segmentation-free, sub-cellular clustering on imaging-based platforms such as Xenium, yet remaining scalable to large Slide-seq, Visium, DSP, and ISS datasets without platform-specific heuristics (**Fig. 1c**).

### Spatial variation uncovers structured gene expression modules in single-nucleus spatial transcriptomic data

We first applied Glimmer to the Slide-tags melanoma dataset^6^, a recently published high-resolution single-nucleus spatial transcriptomics dataset comprising 2,535 cells. Using published cell type annotations only based on transcriptomic features as reference labels, we systematically visualized how tuning the spatial information weight via the log-barrier term in Glimmer shapes the structure of latent embedding. UMAP (Uniform Manifold Approximation and Projection)^36^ shows that when spatial information is excluded, cells are primarily grouped by transcriptional similarity, aligning with published labels (**Fig. 2a, Supplementary Fig. 1a top row**). However, as the weight of spatial information increases, cells of the same type gradually separate into spatially distinct subclusters. At higher weights, two dominant spatial domains emerge, preserving both molecular identity and spatial coherence. This effect is also evident when cells are colored by their spatial coordinates, where the embedding becomes progressively more spatially organized (**Fig. 2b, Supplementary Fig. 1a bottom row**).

**Figure 2.**
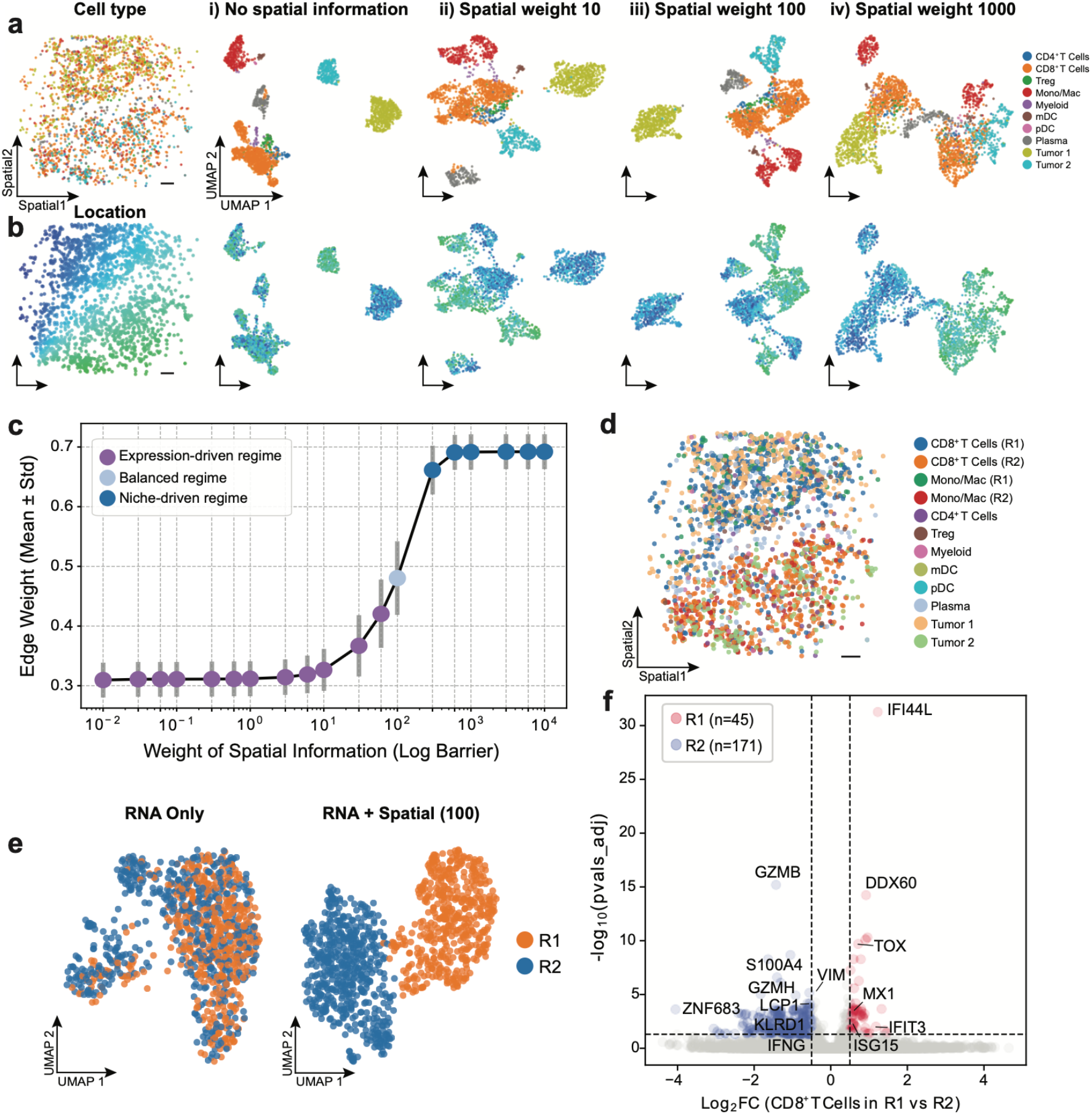
Glimmer reveals biological structure via adaptive balancing of spatial and molecular information in the Slide-tags melanoma dataset. **(a)** Spatial distribution of cell type annotations (left) and UMAPs of RNA-only (i) and Glimmer embeddings with increasing spatial weights (10, 100, 1000) (ii-iv). Higher spatial weights improve the coherence of spatially organized regions. **(b)** Same UMAPs as in (a), colored by spatial coordinates to show spatial alignment across spatial weights. **(c)** Mean ± SD of edge weights in the learned graph across spatial weights. Increasing spatial influence transitions the embedding from expression-driven to spatially organized patterns. Three regimes are defined: expression-driven (low spatial weight), balanced (intermediate), and niche-driven (high). **(d)** Spatial distribution of Glimmer-refined subtypes revealing regional stratification (e.g., CD8^+^ T cells and Mono/Mac) at spatial weight = 100. (e)UMAPs of CD8^+^ T cells under RNA-only (left) and integrated embeddings (right, RNA + spatial, weight = 100), colored by unsupervised clusters from the right (R1/R2, aligned with tumor-dominant regions). Spatial regularization reveals region-specific clusters not captured by transcriptomic data alone. **(f)** Volcano plot showing differential gene expression in CD8^+^ T cells between regions R1 and R2. R1 is enriched for exhaustion markers (e.g., *TOX*), whereas R2 displays elevated cytotoxic genes (e.g., *GZMB, GZMH*). Genes were considered significant at |log_2_FC| > 0.5 and FDR < 0.05. All scale bars: 500 μm.

Notably, BANKSY also uses a tunable parameter λ to control neighborhood influence, similar to Glimmer’s log-barrier. For a fair comparison, we tested multiple λ values on the same dataset and visualized the resulting embeddings. While spatial domain structures were preserved, the embeddings lacked transcriptional coherence, resulting in fragmented or mixed clusters (**Supplementary Fig. 1b**). In contrast, Glimmer achieved a smooth continuum from expression-driven to spatially coherent embeddings, preserving both spatial organization and cell identity. Methodologically, BANKSY augments features by applying a fixed kernel to average neighboring expression values, enforcing local continuity but failing to account for heterogeneity in neighbor contributions. Glimmer instead propagates features over adaptively weighted neighbor graphs, enabling context-aware spatial integration that better captures neighborhood-specific effects while preserving biological identity.

To assess the functional role of the log-barrier in regulating spatial influence, we computed the mean edge weights across all cells under varying parameter settings (**Fig. 2c and Supplementary Fig. 1c**). The weights undergo a sigmoidal transition from an expression-driven regime to a balanced regime and ultimately to a niche-driven regime. The intermediate balanced regime represents an adaptive fusion of spatial and molecular signals, producing spatially informative embeddings while retaining the single-cell nature of the data. Although the precise transition point may vary with platform resolution and tissue context, it can be reliably identified through hyperparameter sweeping or threshold-based tuning tailored to downstream tasks.

We next evaluated whether the balanced regime captures biologically meaningful structure by analyzing a representative setting (spatial weight = 100), examining how the integration of spatial and molecular signals shapes cell organization in the latent space. At this setting, CD8^+^ T cells and Monocyte/Macrophage populations were further subdivided into two transcriptomic states (R1 and R2), whose spatial distributions were distinct and aligned with tumor compartmentalization (**Fig. 2d, Supplementary Fig. 2a**,**b**). This result indicates niche-associated immune remodeling. Within the CD8^+^ T cell subset, RNA-only embeddings failed to distinguish R1 and R2, whereas Glimmer’s spatially informed embeddings resolved them into transcriptomically distinct clusters with spatial segregation (**Fig. 2e**). This underscores the critical role of spatial context in revealing niche-associated immune states. Differential expression analysis showed that CD8^+^ T cells in the R1 state expressed higher levels of exhaustion-related or interferon-stimulated genes such as *TOX, DDX60*, and *IFI44L*^*37,38*^, whereas cells in the R2 state upregulated cytotoxic effectors including *GZMB, GZMH*^*39*^ (**Fig. 2f**). These transcriptomic differences were corroborated by functional scoring, which showed cytotoxicity enrichment in R2 and exhaustion signatures in R1 (**Supplementary Fig. 2c**), indicating functional divergence between spatially defined immune niches.

Glimmer embeddings were further applied to the Monocyte/Macrophage populations to determine whether similar spatially structured heterogeneity could be captured. Consistent with findings in CD8^+^ T cells, the integrated embeddings revealed regionally restricted clusters that were not captured by transcriptomic data alone (**Supplementary Fig. 2a**,**b**). Gene module analysis using Hotspot^40^ in the Glimmer embedding space further identified spatially coherent gene expression programs (**Supplementary Fig. 2e**), which were largely absent when using RNA-only or spatial-only features. Two of these modules (Int Module 14 and 17) exhibited high spatial autocorrelation and corresponded to myeloid programs that are aligned with the functional separation observed in T cells. Modules 14 and 17 showed distinct spatial patterns, with Module 14 enriched in macrophages across Region 1 and colocalized with the Tumor transcriptomic cluster 1 and Module 17 in R2 cells within Tumor transcriptomic cluster 2 areas (**Supplementary Fig. 2f**). Functional analysis (**Supplementary Fig. 2g**) showed that Module 14 was associated with complement and inflammatory pathways, consistent with tumor-associated macrophage activity^41^, while Module 17 was enriched for interferon responses, apoptosis, and *JAK/STAT* signaling, reflecting a more immune-activated state^42^. Together, these findings highlight that balancing spatial and molecular information enables the discovery of niche-specific cellular states that remain obscured in unimodal analyses.

### Glimmer achieves consistent spatial region identification across scales and modalities

We further employ Glimmer to resolve the spatial tissue architecture of a mouse cerebellum section profiled by Slide-seq V2^43^, which captured the expression of 23,096 transcripts across 9,985 spatial beads. Cell type annotations inferred by RCTD^44^ were used as a reference for visualization (**Fig. 3a**). Based on the learned latent representation, Glimmer stratified the tissue into four principal domains corresponding to the canonical laminar organization^45,46^ of the cerebellum: the oligodendrocyte, granule, Purkinje–Bergmann, and molecular layers (**Fig. 3b**). From inner to outer layers, the oligodendrocyte layer (OL) is dominated by oligodendrocytes, the granule layer (GL) consists almost entirely of granule cells, the Purkinje–Bergmann layer (PL) is enriched with Bergmann glia and Purkinje neurons, and the molecular layer (ML) contains a high density of MLI1 and MLI2 interneurons (**Supplementary Fig. 3a**). Layer-specific marker genes from dominant cell types further validate these structural assignments (**Fig. 3c**). For example, *Neurod1* in the GL^47^, *Calb1* in the PL^48^, *Plp1* in the OL^49^, and *Nxph1* in the ML^50^ are consistent with expected cell type distributions.

**Figure 3.**
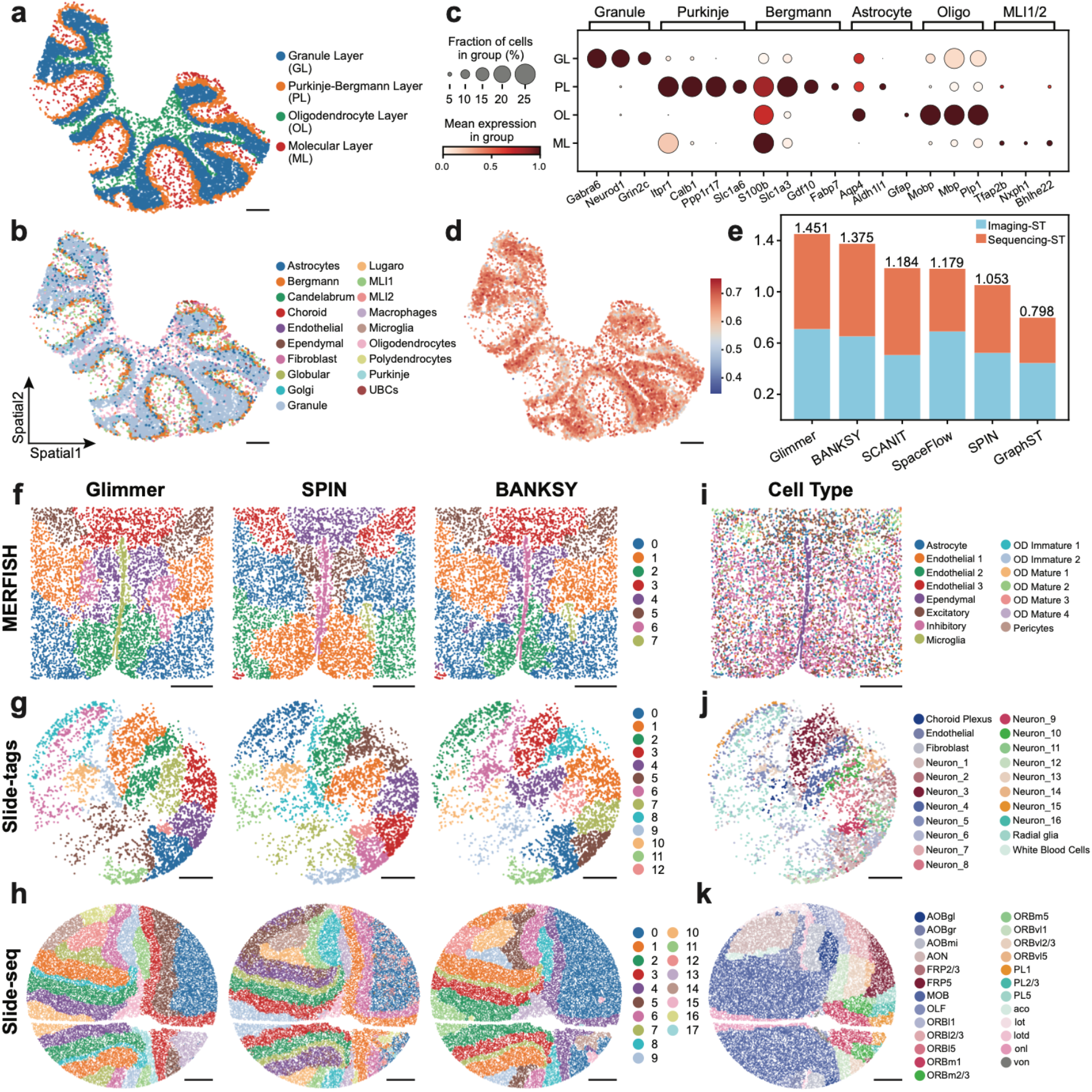
Multi-platform benchmarking of Glimmer demonstrates robust domain identification across diverse spatial transcriptomics platforms. **(a)** Spatial domains in the cerebellum identified by Glimmer, annotated by anatomical layers. **(b)** RCTD-based cell type annotations for comparison. **(c)** Dot plot of canonical cell type markers across cerebellar layers. Dot size indicates the proportion of cells expressing the marker; color intensity reflects average expression level. **(d)** Spatial distribution of mean neighbor weights inferred by Glimmer for each cell. **(e)** Benchmarking performance across MERFISH and Slide-seq mouse brain datasets. Bar height represents the total score (normalized within each platform to [0–1]), with contributions separated by imaging-based and sequencing-based evaluation metrics. **(f-h)** Spatial domain identification results on MERFISH (f), Slide-tags (g), and Slide-seq (h) datasets, as predicted by Glimmer, SPIN, and BANKSY. **(i-k)** Corresponding reference cell type annotations for the datasets shown in (f–h). Scale bars: a, b, d, 250 μm; f–k, 500 μm.

The spatial distribution of mean neighbor weights illustrates how Glimmer leverages local affinities to perform structure-aware smoothing (**Fig. 3d**). To connect this pattern to biological structure, we analyzed the relationship between entropy of cluster proportion and the standard deviation of neighbor weights. Homogeneous regions like the GL exhibit high polarization and mean weights, along with low weight variance, reflecting uniform neighbor contributions. In contrast, the narrow PL shows higher weight variability, suggesting selective neighbor weighting to prevent over-smoothing (**Supplementary Fig. 3b**,**c**).

Next, we evaluated Glimmer alongside existing spatial embedding methods including BANKSY, SCAN-IT, SPIN, SpaceFlow, and GraphST on five region-annotated MERFISH datasets and five unlabeled Slide-seq datasets^51^. For MERFISH datasets, Glimmer achieved the highest adjusted rand index (ARI) and normalized mutual information (NMI) scores and performed strongly on completeness and homogeneity (**Supplementary Fig. 4a**). For Slide-seq datasets, it outperformed other methods on spatial conservation metrics including cell type affinity similarity (CAS) and maximum leiden adjusted mutual information (MLAMI), and achieved comparable performance on niche coherence metrics including niche average silhouette width (NASW), and cell type normalized mutual information (CNMI) (**Supplementary Fig. 4b**). Overall, Glimmer achieved the highest aggregated score and demonstrated strong consistency across sequencing technologies (**Fig. 3e**). Cross-platform regional visualizations using MERFISH, Slide-tags, and Slide-seq (**Fig. 3f–h; Supplementary Fig. 4c–e**) demonstrated that Glimmer consistently identifies spatially coherent and biologically meaningful domains. The identified domains showed strong concordance with reference cell type annotations (**Fig. 3i–k**) and region-level ground truth from previous datasets (**Supplementary Fig. 4f**). Notably, kernel-based frameworks such as Glimmer, SPIN, and BANKSY offer lightweight designs suitable for broad application in spatial analysis.

Beyond transcriptomics, we applied Glimmer to a CODEX protein-based dataset^3^ comprising 8 mouse spleen sections from a systemic lupus erythematosus model spanning two disease stages. By identifying both shared and condition-specific regions across slides, we demonstrated that Glimmer is not only modality-agnostic but also well-suited for cross-sample integrative spatial analysis (**Supplementary Fig. 5a**,**b**).

### Neighbor-guided transcript smoothing enables robust segmentation-free cellular identification in spatial omics

While imaging-based spatial platforms like Xenium provide subcellular-resolution expression and nuclear imaging, they often exhibit mismatches between segmentation masks and transcript distributions^4^. Misassigned transcripts due to inaccurate nuclear boundaries or segmentation artifacts can compromise clustering and differential expression analysis. To address this limitation, we adapted Glimmer to enable segmentation-free cellular clustering directly at the transcript level. Specifically, we applied its neighbor-smoothing framework within a local radius (∼30–50 µm) to approximate the spatial extent of individual cells. As a case study, we selected a patch from Xenium lymph node tissue, discretized transcripts into mini bins (∼2– 3 µm), and reconstructed a bin-level expression matrix (**Fig. 4a**). Under the principle that transcripts belonging to the same cell should exhibit greater expression similarity, we applied neighbor-guided smoothing with a reduced spatial weight to prioritize local transcript coherence. The resulting latent representation enabled bin-level clustering for transcript assignment independent of predefined segmentation masks (**Fig. 4b**). These molecular clusters were subsequently used to reassign transcripts that were incorrectly grouped within a segmented cell, thereby correcting mismatches between morphology and gene expression (**Fig. 4c**).

**Figure 4.**
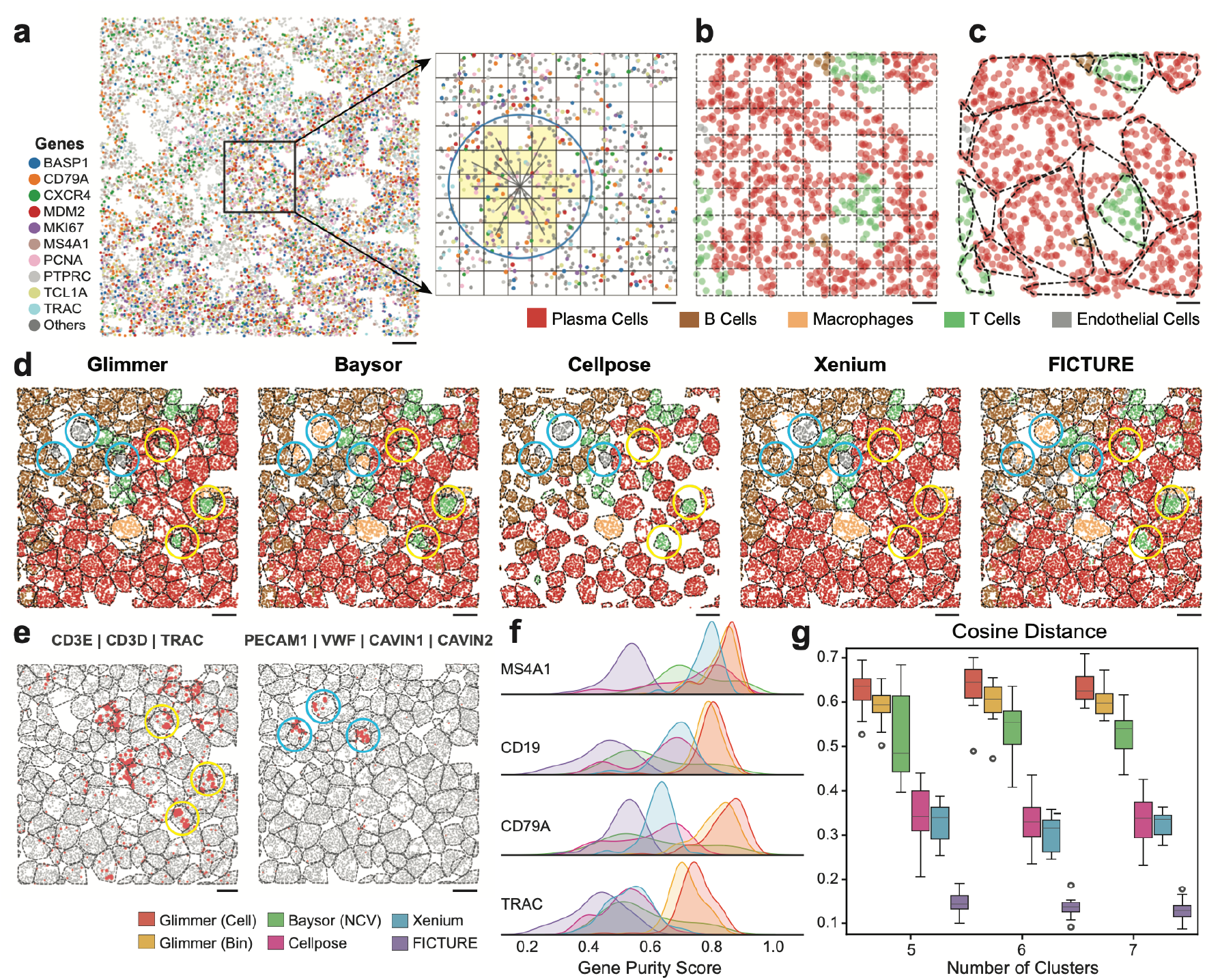
Glimmer enables accurate segmentation-free transcript assignment in imaging ST. **(a)** Spatial locations of selected gene transcripts (e.g., CD79A, TRAC) in a Xenium lymph node patch (left). Zoom-in schematic illustrates Glimmer’s bin-based graph, where each bin aggregates transcript counts and neighboring bins (yellow) are used for graph weight learning and feature smoothing (right). **(b)** Transcripts colored by predicted cellular clusters derived from neighbor-smoothed bin embeddings and annotated by canonical marker genes. **(c)** Voronoi-based segmentation derived from DAPI-stained nuclei and refined by cluster labels from (b), providing cell boundaries for evaluation. **(d)** Cluster assignments predicted by Glimmer, Baysor, Cellpose, Xenium-native segmentation, and FICTURE, with Leiden clustering applied to ensure a consistent number of clusters. Colored by cluster identity. Circles highlight example cells where endothelial cells (blue) and T cells (yellow) are misassigned by other methods. **(e)** Spatial visualization of cell-type–specific markers overlaid with Xenium-defined cell boundaries. The left panel shows T cell markers (*CD3E, CD3D, TRAC*), and the right panel shows endothelial markers (*PECAM1, VWF, CAV1N1, CAV1N2*). Dot size reflects local transcript density smoothed over a 15-neighbor kernel, with larger dots indicating higher local concentration. **(f)** Gene purity scores across methods, defined as the highest proportion of a marker gene’s transcripts assigned to a single cluster (closer to 1 indicates higher cluster specificity). Benchmarking genes include *MS4A1, CD19, CD79A* (B cells), and *TRAC* (T cells). **(g)** Cosine distance between cluster-averaged gene expression profiles across 15 Xenium patches (800 µm × 1600 µm), evaluated at cluster numbers of 5, 6, and 7 over 5 random seeds. Higher distances indicate greater inter-cluster separation. Box plots display the median (center line), interquartile range (box), and 1.5× IQR (whiskers), with outliers shown as individual points. Scale bars: a (left), 10 μm; a (right) and b–e, 2 μm.

To evaluate clustering assignment fidelity, we compared Glimmer against four state-of-the-art segmentation or segmentation-free methods: Baysor^27^, Cellpose^52^, Xenium’s default segmentation, and FICTURE^29^. While all approaches produced broadly consistent transcript-level cluster maps, Glimmer demonstrated superior accuracy in resolving ambiguous transcripts, particularly within minority cell populations in local patches, such as T cells and endothelial cells (**Fig. 4d**). In addition, when examining the spatial expression of canonical marker genes such as *CD3E, CD3D*, and *TRAC* for T cells, and *PECAM1, VWF, CAVIN1*, and *CAVIN2* for endothelial cells^53^, Glimmer’s cluster assignments aligned most closely with known biological patterns (**Fig. 4e**). In contrast, due to the high expression of B cell associated transcripts in the surrounding area, other methods misclassified some cells as B cells, even though very few B cell specific transcripts were actually detected within the corresponding cellular regions (**Supplementary Fig. 6a**). We further compared the marker gene expression profiles across predicted cell types from each method (**Supplementary Fig. 6d**). Glimmer consistently exhibited cleaner expression patterns, characterized by higher expression levels within canonical marker genes for each cell type. For quantitative evaluation, Glimmer significantly outperformed other methods at both cell-level and bin-level clustering, achieving higher transcript assignment purity within target clusters and greater separation between clusters, as measured by marker gene purity (**Fig. 4f**) and cluster distance metrics (**Fig. 4g; Supplementary Fig. 6c**), and detailed in Supplementary Tables 3 and 4.

We further evaluated Glimmer by appling it to two Xenium sections representing follicular lymphoid hyperplasia (FLH) and reactive follicular hyperplasia (RFH) in human tonsils (**Supplementary Fig. 7a–b**). Following batch correction with scVI^54^, Glimmer identified consistent cell type clusters across conditions (**Supplementary Fig. 7a**,**f**), with spatial distributions that aligned with known tonsillar architecture, including germinal centers, mantle zones, and subepithelial stroma (**Supplementary Fig. 7g–h**). Cell type predictions were validated by marker gene expression (**Supplementary Fig. 7e**) and showed compartment-specific enrichment, such as proliferative B cells in germinal centers, CD4^+^ T cells in paracortical zones, and fibroblasts in subepithelial stroma (**Supplementary Fig. 7j**). Notably, Glimmer revealed both shared and distinct spatial patterns between FLH and RFH, highlighting shifts in immune composition such as plasma cell expansion and altered localization of T cells (**Supplementary Fig. 7f, j**). These results demonstrate Glimmer’s capacity to perform integrative, segmentation-free spatial analysis across disease states, capturing cell- and region-level heterogeneity with high resolution.

### Glimmer maps spatial zonation and niche communication in the tonsil microenvironment

The immune system exhibits diverse functional zonation and complex interactions within its microenvironment. To assess whether Glimmer can reveal niche-level functional heterogeneity and intercellular signaling, we applied it to a human tonsil section from the Slide-tags dataset^6^ (5,778 cells, 26,099 genes). Glimmer identified five spatial domains that closely correspond to established anatomical compartments: paracortex, interfollicular zone, dark zone (DZ), light zone (LZ), and marginal zone (MZ), consistent with distribution of published reference cell types^6^ (**Fig. 5a,b**). The spatial expression of germinal center (GC) markers further supported this compartmentalization, with LZ B cell markers (*CD83, CXCL13*) and DZ B cell markers (*AICDA, CXCR4*) displaying mutually exclusive, spatially confined patterns^55^ (**Fig. 5c**). Differential expression analysis between LZ and DZ B cells reinforced these findings (**Fig. 5d**). LZ B cells significantly upregulated *CD74, CD83*, and *CXCL13*, highlighting their roles in antigen presentation and interaction with T follicular helper cells. In contrast, DZ B cells expressed *AICDA, CXCR4, EZH2*, and *FOXO1*, reflecting their involvement in proliferation and somatic hypermutation^55^. While cell type distribution alone makes it difficult to demarcate the LZ and DZ, Glimmer was able to detect nuanced spatial patterns and distinguish these biologically distinct compartments.

**Figure 5.**
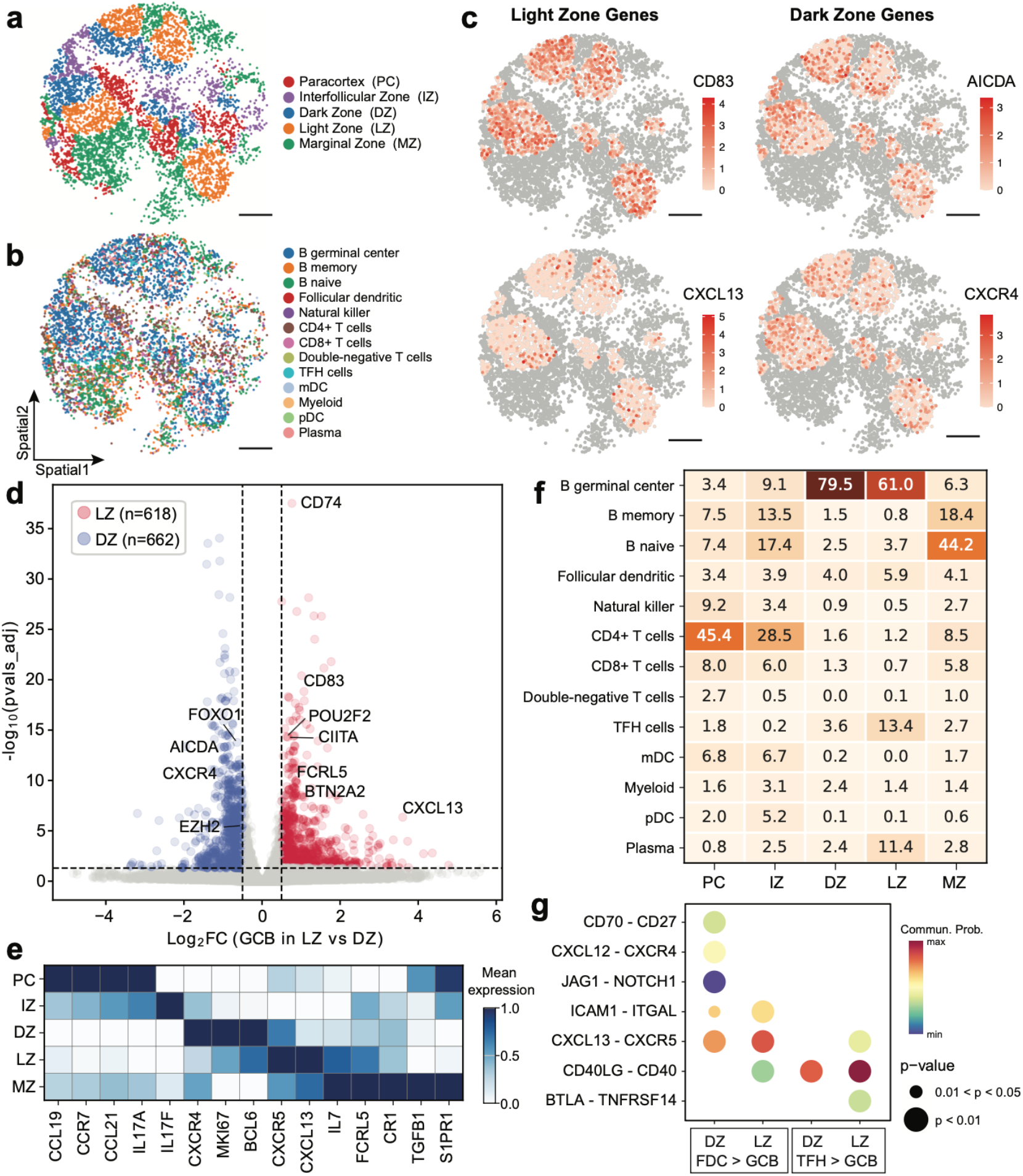
Glimmer’s enables in-depth dissection of niche-specific immune signaling in the human tonsil. **(a)** Spatial regions (R1–R5) identified by Glimmer, annotated with anatomical zones (top) and corresponding cell type annotations based on published references (bottom). **(b)** Spatial expression of canonical germinal center B cell markers distinguishing light zone (LZ; *CD83, CXCL13*) and dark zone (DZ; *AICDA, CXCR4*). **(c)** Volcano plot showing differentially expressed genes between LZ and DZ germinal center B cells. X-axis shows log_2_ fold change (LZ vs. DZ); Y-axis shows –log_10_ adjusted p-values. Genes with |log_2_FC| > 0.5 and FDR < 0.05 are considered significant. **(d)** Heatmap showing the expression patterns of key ligands and cytokines involved in orchestrating B cell development across regions. **(e)** Distribution (%) of immune cell types across Glimmer-defined spatial regions. **(f)** Region-specific ligand–receptor interactions between follicular dendritic cells (FDCs) and germinal center B cells inferred from CellChat. Dot size indicates statistical significance; color denotes communication probability. All scale bars: 500 μm.

Spatial immune regions are often occupied by specific cell–cell interactions, where ligands and cytokines orchestrate these signalings. To investigate molecular signals specialization across spatial niches, We examined the expression of key ligands, receptors, cytokines, and regulatory genes within these regions. The heatmap reveals spatial gradients of molecules critical for B cell migration and GC organization (**Fig. 5e**). For example, *CXCL13* and *IL7* were enriched in the LZ and MZ, guiding B cell recruitment into LZ and surrounding follicular regions. *CXCR5*, the receptor for *CXCL13*, also peaked in the LZ, consistent with its role in retaining B cells within GC^56^. In contrast, *S1PR1*, which promotes B cell egress, was elevated in the MZ, supporting a directional model of B cell maturation and exit from the follicle^57^. Together, these findings highlight a spatially organized cytokine milieu that coordinates B cell trafficking during affinity maturation.

After characterizing gene expression, we next assessed cell-type composition across domains to uncover tissue-level organization (**Fig. 5f**). GC B cells localized to DZ and LZ, while CD4^+^ T and T follicular helper (TFH) cells concentrated in the paracortex and interfollicular zone, consistent with their helper roles and known lymphoid compartmentalization. To explore region-specific cellular interactions, we applied CellChat^58^ to infer ligand–receptor signaling networks within LZ and DZ separately (**Fig. 5g**). Signaling from follicular dendritic cells to B cells was enriched in the DZ and LZ, with *CXCL12-CXCR4* and *CXCL13-CXCR5* as key pairs supporting B cell survival and positioning^59^. TFH–B cell communication in the LZ was dominated by *CD40LG-CD40* and *BTLA-TNFRSF14*, mediating costimulatory and synapse-forming interactions^60,61^. These ligand–receptor pairs were spatially enriched, underscoring zone-specific communication programs that shape B cell fate within GC. These findings highlight Glimmer’s ability to decode zone-specific communication and spatial architecture, offering a unified framework for microenvironmental analysis in complex tissues.

## Discussion

Biologically meaningful spatial representations underpin a broad range of downstream tasks in spatial omics, from cell type annotation to niche discovery and integrative clustering across samples. In spatial transcriptomics data, spatial coordinates reflect physical context and potential interactions, while gene expression encodes functional identity. Existing methods rarely model this coordination explicitly; they rely on static neighborhood graphs or task-specific heuristics. These hard-coded choices often break when tissue architecture, resolution, or omics modality, producing embeddings that either over-smooth informative biological variation or fragment spatially and molecularly coherent regions.

Glimmer reframes representation learning as neighbourhood-graph optimisation. By minimising Dirichlet energy, it iteratively rewires edge weights so that spatially proximate, expression-similar cells reinforce one another, while preserving all local connections with adaptively learned contributions. A distance-based decay limits the influence of distant neighbours, and a log-barrier term prevents degenerate weights, while bounded soft weights promote stability and adaptively scale each neighbor’s influence. Together, these mechanisms enable the learning of context-aware graphs that capture biologically meaningful structure from sub-cellular bins to millimetre-scale regions. Crucially, it makes neighbor selection and weighting transparent, allowing tracing of how embeddings arise from local neighborhood context.

The resulting embeddings preserve biologically informative gradients, while enabling local smoothing of spatial data, providing a versatile approach for tasks ranging from tissue domain segmentation to cellular segmentation of imaging spatial transcriptomics. In the human tonsil, Glimmer recovers known regions and reveals functional niches shaped by cell interactions and gene gradients. In imaging-based platforms like Xenium, it improves cell clustering precision by leveraging local spatial coherence without relying on explicit cell boundaries. Moveover, Glimmer’s local graph structure also supports cross-sample integration, enabling comparison of shared and condition-specific programs across tissues and states.

Despite its strengths, Glimmer has several limitations that offer promising avenues for future work. First, although most parameters – such as sparsity and neighborhood radius – tend to be robust across datasets, the log-barrier weight may require dataset-specific adjustment. Developing generalizable heuristics from large-scale benchmarks, or incorporating automated, data-driven strategies for hyperparameter selection, could further improve the robustness and usability of the framework. Second, although the current implementation based on first-order graphs performs well, Glimmer’s underlying principle naturally extends to higher-order graphs. While first-order graphs offer a good balance between performance and efficiency, second-order graphs could potentially capture more complex neighborhood dependencies. Finally, although Glimmer has been demonstrated primarily on transcriptomics data, the graph-based formulation is modality-agnostic. Future work may explore the integration of multi-omic spatial signals and even temporal extensions for modeling tissue evolution during development, regeneration, or disease progression.

In summary, Glimmer combines a conceptually rigorous graph optimization approach with biological interpretability to provide a versatile toolkit for learning spatially informed representations from spatial omics data. It accurately assigns transcripts to clusters without relying on segmentation, enabling subcellular-resolution analysis. The method captures spatial architecture, reveals cellular niches, and supports integrative clustering across samples, while remaining compatible with various omics modalities. We anticipate that this framework will serve as a foundation for exploring cellular networks and spatial dynamics in increasingly complex biological systems.

## Methods

### Model Formulation

In biological systems, the underlying graph structure that encodes cellular relationships is not directly observable and must be inferred from the data. Assuming that cellular features vary smoothly over a meaningful latent graph, we formulate the problem of learning edge weights as minimization of Dirichlet energy which serves as a principled metric measuring the local continuity reflected by the structure. This approach facilitates the construction of biologically informed neighborhood graphs that integrate both feature similarity and spatial proximity.

Let *G* = (*V, E, W*) be a weighted undirected graph, where *V* is the set of nodes, *E* is the set of edges, and *W* is the symmetric weight matrix with non-negative entries *w*_*ij*_ ≥ o. The Dirichlet energy of a function *f*: *V* → *R*^*d*^ (*d* is the dimensionality of the embedding space) is given by:

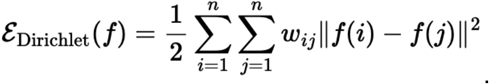

This energy measures the smoothness of *f* over the graph and is minimized when neighboring nodes have similar representations. It plays a central role in many geometric learning methods, such as Laplacian eigenmaps, spectral clustering, and graph-based regularization^62–64^.

In our framework, we extend the Dirichlet energy formulation by introducing biologically meaningful spatial constraints through an edge-weighting scheme. Specifically, we 1) apply an exponential decay to down-weight edges based on physical proximity, introducing distance-aware local influence; 2) transform the decayed weights using a sigmoid activation to ensure boundedness and numerical stability; and 3) incorporate two regularization terms—a logarithmic barrier that prevents degenerate solutions and encourages distributed edge connectivity, and an *l*_2_ norm that controls the overall weight magnitude and stabilizes optimization^34^. These combined modifications result in a spatial graph that captures smooth functional transitions and underlying tissue organization.

We begin by defining the mathematical components of the spatial graph. Let *X* ∈ *R*^*n*×*m*^ be the feature matrix, where *n* is the number of cells and *m* the number of feature dimensions (e.g., PCA-reduced expression profiles). For each cell *i*, we construct a first-order spatial graph by identifying its *k*-nearest neighbors based on Euclidean distances in physical space. The resulting distance matrix *S* ∈ *R*^*n*×*k*^, computed as the inverse of the spatial distance between locations, is then scaled to the range [0, 1] and used as a spatial decay factor to modulate the strength of local interactions. To quantify feature similarity, we define the pairwise feature distance matrix *Z* ∈ *R*^*n*×*k*^ where each entry is given by:

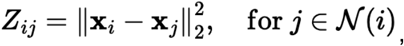

where *N*(*i*) denotes the set of *k*-nearest neighbors of cell *i*.

Let *W* ∈ *R*^*n*×*k*^ denote the edge weight matrix, which is the primary optimization variable and learned by minimizing a regularized Dirichlet energy objective:

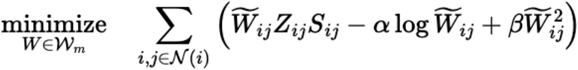

with

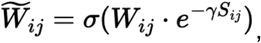

where *σ*(*x*) = 1/(1 + *e*^−*x*^) is the sigmoid activation function that normalizes edge weights into the [0,1] range.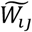, denotes the spatially decayed and sigmoid-activated edge weight. The term 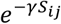 softly attenuates edges between physically adjacent cells, helping to reduce overemphasis on local proximity and promote the discovery of functionally informative distal interactions. The hyperparameters in the objective provide appropriate regularization: *α* > 0 prevents degenerate (i.e., trivial or all-zero) weights, and it is important for balancing the effects of feature similarity and spatial proximity; *β* > 0 controls the degree of sparsity via *l*_2_regularization; and *γ* ≥ 0 controls the sharpness of spatial decay.

After optimization, the final embeddings of cells are computed using the learned graph structure. Specifically, for each cell, its final embedding is computed by aggregating the initial embeddings of itself and its neighbored cells:

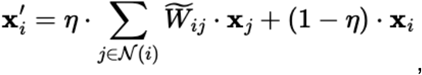

where *η* ∈ [0, 1] determines the strength of neighbor smoothing.

### Training Details

We initialized *W* from a standard normal distribution *N*(0, 1), and optimized it using the Adam optimizer^65^ with a learning rate of 1 × 10^−4^ for 10,000 epochs and a fixed random seed of 42. We used *k* = 50 for niche-driven or spatial regional clustering, and *k* = 15 for expression-driven subcellular clustering. The setting of the hyperparameter *α* depends on specific technology and tissue types. In general, we recommend using a large *α* when 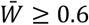 for regional clustering, and a smaller *α* when 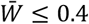 for subcellular clustering, where 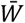 denotes the mean of all elements in the trained weight matrix *W*. We used *α* = 1,000 as the initial value. The hyperparameters *β* and *γ* were fixed at 0.01 and 1, respectively. The feature smoothing interpolation parameter *η* was set to 0.1 for regional clustering and 0.2 for subcellular clustering. All models were trained on a single NVIDIA GPU (RTX A6000) using PyTorch 2.7.0 with CUDA 12.6 support.

### Benchmarking

#### (a) Niche-driven clustering for domain identification

##### Datasets and processing

We curated two types of datasets for benchmarking. The first consists of 5 region-annotated MERFISH slices from the mouse hypothalamic preoptic region^1^, each containing 5,488 to 5,926 cells and sharing a common set of 155 genes. The second includes 5 Slide-seq samples from coronal sections of the mouse brain atlas^46^. Only cells with valid anatomical annotations were retained. To reduce computational overhead and standardize cell numbers, cells within a circular region centered at the geometric mean of spatial coordinates were selected, with the radius adjusted to yield 30,000–40,000 cells per sample. All datasets were preprocessed using Scanpy (version 1.10.4). Gene expression counts were normalized to 10,000 per cell and log-transformed. Up to 3,000 highly variable genes were selected using the Seurat v3 method. PCA was performed with the ARPACK solver, and the top 50 components were retained for downstream analyses.

##### Evaluation metrics

For datasets with ground-truth regional annotations, we evaluated clustering performance using Adjusted Rand Index (ARI), Normalized Mutual Information (NMI), completeness, and homogeneity^51^. ARI and NMI quantify the agreement between predicted and true cluster assignments, while completeness and homogeneity assess whether all samples from the same class are assigned to the same cluster and whether each cluster contains only samples from a single class, respectively. All metrics range from 0 to 1, with higher values indicating better clustering quality. For Slide-seq datasets without well-defined region labels, we used Cell Type Affinity Similarity (CAS) and Maximum Leiden Adjusted Mutual Information (MLAMI)^66^ to evaluate the spatial conservation of latent representations. CAS compares cell-type enrichment in local neighborhoods between latent and physical spaces, while MLAMI quantifies the overlap between clusters derived from latent embeddings and those obtained based on spatial proximity. We also computed Niche Average Silhouette Width (NASW) and Cell Type Normalized Mutual Information (CNMI) to assess the structural coherence of the latent space. NASW was calculated by applying Leiden clustering at resolutions and averaging the silhouette widths across clusterings. CNMI was computed following the implementation provided in scIB (v1.1.7)^67^.

##### Implementation details

We compared Glimmer with five state-of-the-art spatial clustering methods including GraphST (v1.1.1), SpaceFlow (v1.0.4), SCAN-IT (v0.1), SPIN (v0.0.1), and BANKSY (v0.99.12). All models were executed using their default or recommended settings. Specifically, GraphST was run with the data type set to *slide mode*. SpaceFlow was trained with a latent dimension of 50 and a spatial regularization strength of 0.1. SCAN-IT was performed with spatial graphs constructed using the *alpha shape* method with 2 layers and 5 neighbors, followed by multidimensional scaling to obtain latent embeddings of dimension 30. BANKSY was implemented in R (v4.3.1) using *normalized counts* as input, with spatial neighborhood sizes of 15 and 30, and a regularization parameter λ of 0.8. Latent embeddings were obtained from PCA-reduced components. For Glimmer, the spatial graph was constructed using 50 spatial nearest neighbors, with a log-barrier weight of 5000 and a neighbor weight of 0.1. Except for BANKSY and SpaceFlow, which used their own preprocessing steps, all methods explicitly used PCA matrices as input. Each method was evaluated using 5 random seeds per dataset. For reproducibility, random seeds for Python, NumPy, and PyTorch were fixed across all runs.

For MERFISH datasets, Leiden clustering was performed using the default 20 nearest neighbors in the output embeddings, with a shared set of resolution values ranging from 0.05 to 0.5 in increments of 0.05. Only clustering results in which the number of clusters fell within a ±3 range of the ground-truth cluster count were retained for evaluation. For Slide-seq, all metrics were computed using 15 nearest neighbors in both the latent space and spatial coordinates, following the published settings. NASW used Leiden resolutions of 0.1, 0.55, and 1.0. The final scores were computed by averaging the metrics across one sample.

#### (b) Expression-driven clustering at subcellular level

##### Datasets and processing

The Xenium slide from a non-diseased human lymph node was partitioned into 15 batches for evaluation. Each batch comprised two adjacent fields of view (FOVs) to preserve transcript capture continuity across FOV boundaries and to evaluate potential model biases toward local capture patterns. All *NegControlProbe* and *BLANK* transcripts were removed prior to benchmarking. For the Glimmer binning-based clustering strategy, we began by randomly selecting 100 transcripts to estimate the average minimal bin diameter required to include at least 5 transcripts, starting the search with a bin size of 1um and a step size of 1. In general, all patches converged to a bin size of 2 μm, yielding bin-level gene expression profiles for downstream clustering.

##### Evaluation metrics

The purpose of benchmarking was to evaluate clustering purity, without assessing segmentation quality. Cosine distance and Jensen-Shannon divergence were used to quantify inter-cluster feature dissimilarity. Gene purity was used to assess the biological validity of clustering results. Specifically, we first computed the average gene expression profile for each cluster under each Leiden or K-means resolution, then calculated pairwise distances between clusters based on these profiles. For each clustering resolution, gene purity was defined as the highest proportion of transcripts for each marker gene (MS4A1, CD19, CD79A, and TRAC), observed across clusters and calculated based on the original transcript counts. Only clustering results with 5, 6, or 7 clusters were considered for benchmarking, reflecting the expected number of major clusters.

##### Implementation details

We evaluated the clustering performance of Glimmer in comparison with segmentation-based approaches including Cellpose3 (v3.0.10) and Xenium (default segmentation), as well as segmentation-free methods including Baysor (v0.7.1, Neighborhood Composition Vector mode, NCV) and FICTURE (v0.0.1). For Cellpose, we subset paired and overlapping DAPI images from the corresponding FOVs within the whole-slide dataset. The model was trained using the *cyto3* backbone with the estimated cell diameter set to 50. For Xenium, the default segmented cell-by-gene expression matrix was directly used for clustering analysis. Leiden clustering was performed for both Cellpose and Xenium on the segmented expression matrices using resolutions from 0.4 to 0.9 with a step size of 0.05. For Baysor, NCV mode was applied with *k* = 15 for K-nearest neighbors, a minimum cell size of 15 molecules, and default 3D spatial coordinates (x, y, z). Each transcript was assigned an RGB color representing its local neighborhood composition. These colors were converted to CIELAB color space, and K-means clustering (*k* = 5, 6, 7) was applied to the LAB values to identify spatial clusters. FICTURE was executed using its all default settings, with the number of clusters set to 5, 6, and 7. For Glimmer, we first constructed a binned transcript expression matrix, where the bin size was determined based on the average diameter required to include at least five transcripts for 100 randomly selected transcripts. The spatial coordinates of each bin were defined as the mean x and y positions of all transcripts within that bin. For the binned spatial datasets, we first applied PCA to reduce dimensionality, followed by construction of a spatial neighbor graph using the 15 nearest spatial neighbors. The graph used a log-barrier weight of 200 and a neighbor weight of 0.2. The resulting embeddings were then subjected to Leiden clustering, with resolution values ranging from 0.4 to 0.9 in steps of 0.05.

### Analysis

#### (1) Slide-tags human melanoma

##### Spatial embedding using Glimmer

Human melanoma spatial transcriptomic data were processed using Scanpy, including normalization, log-transformation, selection of 3,000 highly variable genes (HVGs), and principal component analysis (PCA) on the top 50 components. Glimmer was applied with 50 nearest spatial neighbors, using log-barrier values ranging from 0.01 to 10,000 to balance transcriptomic fidelity and spatial coherence. The neighborhood graph was constructed based on the smoothed embedding with default Scanpy settings. Spatially informed clusters (Fig. 2d) were identified via Leiden clustering (resolution = 0.8) and annotated by cross-referencing with published cell type labels. Major immune populations were further resolved into subclusters R1 and R2, distinguished by their spatial segregation and differential overlap with two tumor subclones.

##### Immune subtype stratification

CD8^+^ T cells and monocytes/macrophages were subsetted and reprocessed with normalization, log-transformation, selection of 3,000 HVGs, and PCA. Glimmer was applied with a log-barrier value of 100 to generate embeddings, followed by Leiden clustering at resolution 0.2. Differentially expressed genes between clusters were identified using *rank_genes_groups*. Cytotoxicity and exhaustion module scores were computed using *score_genes* based on predefined gene sets. The cytotoxicity gene set included *PRF1, GZMB, GZMA, GZMH, GZMK, GZMM, IFNG, TNF, FASLG, GNLY, SH2D1A, RAB27A, KLRD1, NKG7, KLRK1, CD28, ICOS, TBX21, EOMES, RUNX3, ITGAE*. The exhaustion gene set included *TOX, TOX2, NR4A1, NR4A2, BATF, IRF4, PDCD1, HAVCR2, LAG3, TIGIT, CTLA4, CD244, ENTPD1, CD38, LAYN*. Cells expressing ≥5 genes from either signature were retained for further analysis. Module scores were calculated as the mean expression per cell and compared between R1 and R2 using a two-sided Mann–Whitney U test.

##### Gene module discovery with Hotspot

Hotspot (v1.1.3) was applied to identify gene modules based on three types of embedding: RNA-only (PCA), spatial-only (XY coordinates), and Glimmer-integrated (log_barrier = 100). A denoised negative binomial (danb) model was applied, along with an unweighted k-nearest neighbor graph (*k* = 100), to compute local gene autocorrelation. Embedding-specific modularity was assessed by Jaccard similarity between Glimmer-derived modules and those from RNA-only and spatial-only embeddings. The two Glimmer-specific modules with the lowest maximum similarity were selected for spatial visualization and functional annotation using Enrichr (gseapy v1.1.8).

#### (2) Slide-seq mouse cerebellum

##### Spatial region annotation

Standard preprocessing was applied as above. Glimmer embedding was computed with parameters k = 50, log_barrier_w = 120, and neighbor_weight = 0.1. The neighborhood graph was constructed based on the smoothed embedding using Scanpy’s default settings. Leiden clustering was performed at resolution = 0.6. To reduce local noise, cell labels were smoothed by assigning each cell the most frequent label among its k = 10 nearest spatial neighbors. The resulting clusters were used for region annotation.

##### Regional evaluation of neighbor graph structure

To evaluate whether the learned neighbor graph reflects biologically meaningful structure, its relationship to cellular heterogeneity was analyzed across spatially defined regions. Cell type heterogeneity was quantified using Shannon entropy, computed from the normalized proportions of cell types within each region and further normalized by the log-transformed number of observed cell types to account for regional complexity. Regional purity was defined as the highest proportion of any single cell type within a region. Graph consistency was characterized by the mean and scaled standard deviation (×100) of per-cell average edge weights within each region. These metrics together capture how local graph structure aligns with underlying cellular composition.

#### (3) CODEX mouse spleen slides

Eight publicly available mouse spleen sections from CODEX spatial proteomics data with cell type annotations were analyzed. Spatial coordinates were adjusted using a custom offsetting strategy, sequentially shifting each tissue slice in X and Y by the previous slice’s extent plus a fixed offset (1e12), to prevent overlap across samples in the neighbor graph. Glimmer was applied to the adjusted and concatenated tissue sections using the original expression matrix of 29 proteins. The model was trained with 50 nearest neighbors, a log_barrier of 1e30, a neighbor weight of 0.2, and a batch size of 8,192. No batch effect correction was performed prior to embedding learning. After learning the smoothed Glimmer embedding, a neighbor graph was constructed, and Leiden clustering was performed at a resolution of 0.8. To reduce local noise, cell labels were refined by majority voting among each cell’s 10 nearest spatial neighbors. The resulting clusters were used for region visualization.

#### (4) Xenium human lymph node slide

##### Construction of bin-based expression matrix

Transcriptomic data were aggregated into fixed spatial bins by assigning each transcript to a bin and summing gene counts, yielding a bin-level expression matrix for downstream analyses such as cell type mapping and neighborhood correlation. To determine bin size, 100 spatial locations were randomly sampled, and square bins centered at each point were expanded until at least 50 transcripts were captured. The average of these minimal sizes defined the final bin size, adapting resolution to local transcript density. Spatial coordinates were then discretized into bin indices, and each transcript was assigned accordingly. For each bin, center coordinates, total transcript count, and gene-wise expression levels were recorded. The resulting bin-level matrix supports downstream analyses including Glimmer embedding and Leiden clustering, while transcript-to-bin assignments enable propagation of cluster labels to the transcript level.

##### Voronoi segmentation and geometric refinement

A Voronoi-based spatial segmentation framework was implemented to assign transcripts to individual cells by integrating nucleus localization, cellular expansion, clustering refinement, and geometric filtering. Transcript coordinates were converted from microns to pixel indices based on the imaging resolution (0.2125 μm/pixel) and initially mapped to nucleus IDs derived from a DAPI-based segmentation mask. For each nucleus associated with ≥3 transcripts, a convex hull was generated to approximate its cellular boundary, and both the polygon and corresponding centroid were stored in well-known binary (WKB) format. Nuclei with insufficient coverage or invalid geometry were excluded from downstream processing. A global Voronoi tessellation was constructed from the centroids of valid nuclei and clipped to the tissue boundary to define spatial catchment zones. Transcripts lacking initial nucleus assignments were reassigned based on a KDTree query of centroids, conditional on three spatial criteria: inclusion within the Voronoi region of a candidate nucleus, exclusion from the nucleus’s original convex hull (to preserve high-confidence assignments), and proximity within an adaptive threshold determined from the median cell area. To capture transcriptional heterogeneity within individual Voronoi regions, Leiden clustering labels were incorporated. Transcripts within the same spatial region but belonging to different clusters were assigned distinct cell identities using cluster-specific suffixes (e.g., Cell-13-A), enabling subcellular resolution of mixed populations. To eliminate spatial outliers, inter-cell distances were calculated from centroid coordinates using KDTree-based nearest-neighbor analysis. Cells whose centroids exceeded three times the median nearest-neighbor distance were excluded. Fragments with fewer than five UMIs were flagged as low-coverage cells and merged into the nearest larger cell from the same cluster if spatially proximal; fragments that remained unmatched were optionally discarded. Alpha shapes were constructed to identify spatially nested fragments. Cells in the lowest 5th percentile of area were evaluated for geometric containment, and those with ≥95% of their alpha shape enclosed within a nearby larger cell were removed. Polygon containment analyses were parallelized to ensure computational efficiency. This pipeline yielded high confidence, spatially resolved cell assignments, robust to transcriptional heterogeneity and geometric artifacts.

#### (5) Xenium human FLH and RFH

##### Batch effect correction and integration with scVI

Two human tonsil samples were obtained from the 10x Genomics Xenium public dataset, representing Follicular Lymphoid Hyperplasia (FLH) and Reactive Follicular Hyperplasia (RFH). To reduce data volume while retaining biologically relevant regions, spatial subregions were manually selected from each slide. For FLH, fields of view (FOVs) I7–I11, J7–J11, and K7–K11 were included; for RFH, regions W14–W18, X14–X18, and Y14–Y18 were selected. These areas correspond to densely populated lymphoid zones with active transcriptional activity and were used for downstream integrative analysis. Following subsetting, gene expression matrices from both samples were merged and harmonized using the scVI^54^ (scvi-tools v1.0.3), with slide identity encoded as the batch key. A 50-dimensional latent representation was learned using a negative binomial likelihood with gene-batch-specific dispersion, and the model was trained for 500 epochs with GPU acceleration. The resulting embedding was used for joint representation and comparative analysis across disease conditions.

##### Dual-mode clustering based on Glimmer-informed joint embedding

Following scVI-based integration, spatial clustering was performed using the Glimmer framework to jointly model gene expression and spatial context. Two clustering strategies were applied based on spatial granularity: (1) Expression-driven clustering: Transcripts were aggregated into uniform spatial bins to construct a bin-level expression matrix, log-normalized, and combined with spatial coordinates and scVI-derived embeddings. Glimmer was run with parameters k=15, log_barrier_w=200, and neighbor_weight=0.2. Clustering was performed using the Leiden algorithm at resolution 1.0, and bin-level labels were mapped back to individual cells for visualization. (2) Niche-driven clustering: Cell-level expression matrices from Xenium’s default segmentation were used directly. Glimmer parameters were adjusted to capture finer spatial niches: k=50, log_barrier_w=5, and neighbor_weight=0.1. Leiden clustering was applied with resolution = 0.8, enabling identification of localized transcriptional neighborhoods across conditions.

#### (6) Slide-tags human tonsil

##### Spatial region annotation

Standard preprocessing was applied as above. A Glimmer-based embedding was computed with parameters k = 50, log_barrier_w = 1000, and neighbor_weight = 0.1. The resulting smoothed embedding was used to construct a neighborhood graph with Scanpy. Leiden clustering was applied with resolution = 0.3, and cluster labels were spatially smoothed by majority voting among neighboring cells. The final clusters were used for spatial region annotation.

##### Differential gene expression analysis

GC B cells were subset and further filtered to include only cells from the DZ and LZ regions. Differential expression analysis between zones was performed using Scanpy’s rank_genes_groups function with DZ as the reference. Genes with |log_2_FC| > 17 were excluded as outliers. Significance was defined as adjusted p-value < 0.05 and |log_2_FC| > 0.5.

##### Intercellular signaling in the germinal center

Following region annotation, the LZ and DZ were selected for pairwise analysis. Cell-cell communication was inferred using CellChat (v2.1.2), with communication probabilities estimated via a truncated mean model and contact-based filtering (interaction.range = 100 µm, contact.range = 100 µm, scale.distance = 1, contact.dependent = TRUE). Interactions with fewer than two cells per group were excluded. Pathway-level scores were computed, and networks were aggregated across pathways. Communication strength and significance were visualized using ggplot2-based dot plots, with interaction probability mapped to color and p-value category to point size.

## Data availability

All datasets used in this study were obtained from publicly available resources. Original data source and direct download links are provided in **Supplementary Table 1**.

## Code availability

The Glimmer framework and benchmarking code are released as an open-source Python package, available at https://github.com/thechenlab/Glimmer.

## Supplementary Materials

**Supplementary Figures 1–7**.

Extended results referenced in the main text.

**Supplementary Table 1**.

Source information for all datasets used in the study.

**Supplementary Table 2**.

Hotspot gene modules identified with melanoma dataset.

**Supplementary Table 3**.

Benchmarking scores for segmentation-free clustering accuracy.

**Supplementary Table 4**.

Summary statistics of all benchmarking evaluations for segmentation.

## Acknowledgements

We thank Andrew J. C. Russell, Dawn Chen and Sandeep Kambhampati for their helpful discussions and feedback. We also acknowledge support from the Eric and Wendy Schmidt Center at the Broad Institute.

