## Supplementary Figures for "Energy-Regularized Graph Learning for Multiscale Spatial Representation"

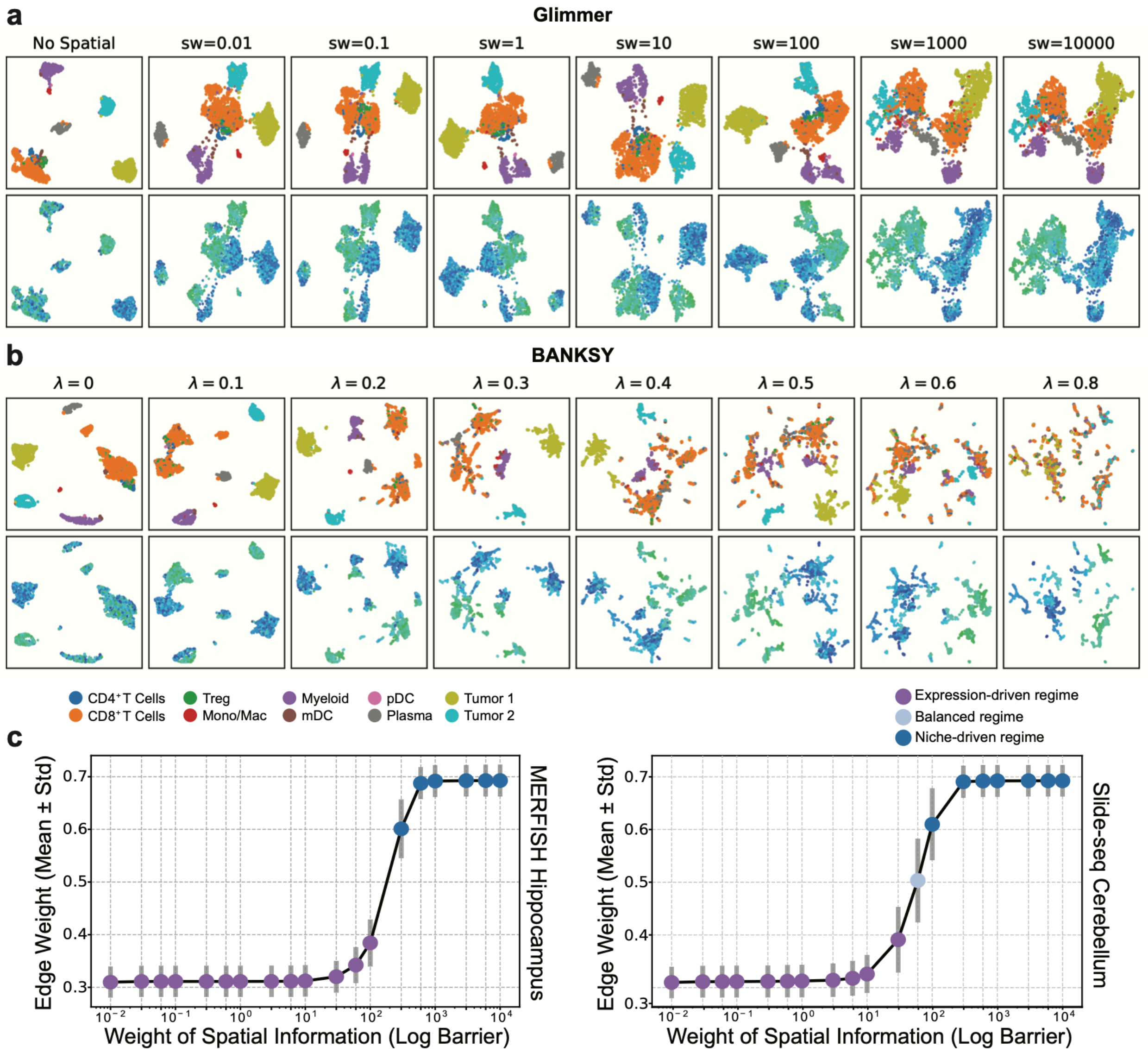


**Supplementary Figure 1 | Robustness of Glimmer embeddings and downstream biological insights.**

**(a–b)** UMAP embeddings from Glimmer (a) and BANKSY (b) across a range of spatial weights (Glimmer: sw = 0.01 to 10,000) and regularization parameters (BANKSY: λ = 0 to 0.8). Top rows are colored by published cell type annotations; bottom rows are colored by spatial coordinates. Glimmer embeddings progressively integrate spatial information while preserving biological continuity. In contrast, BANKSY shows fragmentation and loss of cell-type coherence at higher λ values.

**(c–d)** Parameter robustness of Glimmer under varying spatial information weights (log barrier). Dot plots show edge weight heterogeneity (mean ± std) in MERFISH hippocampus (left) and Slide-seq cerebellum (right), with shaded regimes indicating spatial organization modes.

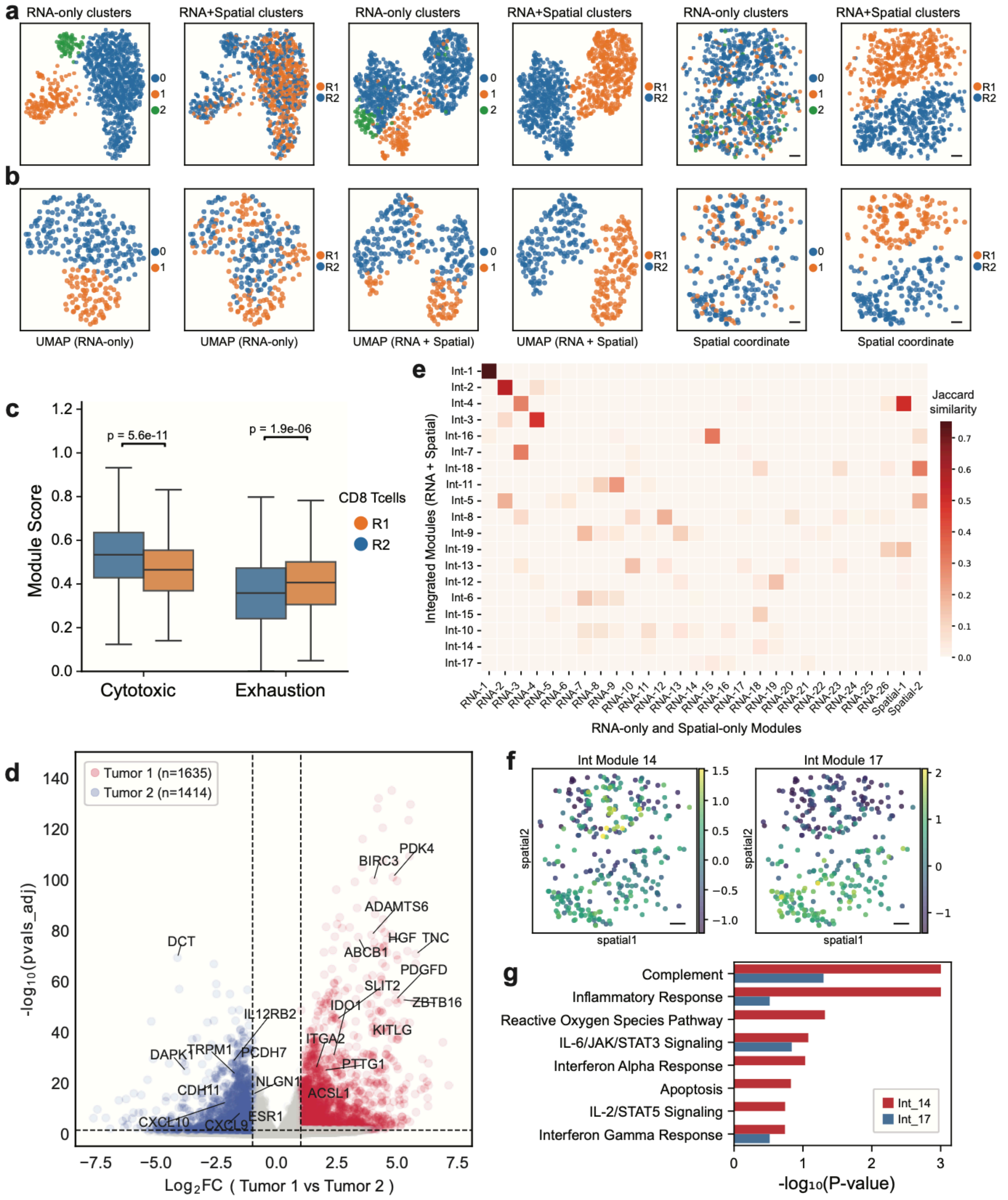

**Supplementary Figure 2 | Glimmer embeddings reveal spatially associated clusters with improved biological interpretability.**
**(a–b)** UMAP of CD8$^+$ T cells (c) and monocyte/macrophage subsets (d). From left to right: UMAPs based on RNA-only embeddings (first two panels), integrated embeddings combining RNA and spatial information (weight = 100; middle two panels), and spatial

coordinates (last two panels). Clustering was performed separately on RNA-only and integrated features, resulting in RNA-only clusters and RNA+Spatial clusters, respectively.
**(c)** Boxplots comparing cytotoxicity and exhaustion module scores of CD8$^+$ T cells between R1 and R2. R2 cells exhibit higher cytotoxic activity, while R1 cells show elevated exhaustion signatures.
**(d)** Volcano plot of differentially expressed genes between Tumor 1 and Tumor 2, spatially corresponding to immune regions R1 and R2. Tumor 1 shows upregulation of genes associated with proliferation, invasion, and immune evasion (e.g., *PTTG1*, *HGF*, *IDO1*), suggesting a more aggressive phenotype. Tumor 2 expresses genes linked to immune activation and differentiation (e.g., *CXCL10*, *DAPK1*, *DCT*). Dashed lines indicate thresholds for significance: $|\log_2 FC| > 1$ and FDR < 0.05.
**(e)** Heatmap showing Jaccard similarity between modules derived from integrated embeddings (RNA + spatial, weight = 100, Y axis) and those from RNA-only features or spatial coordinates (X axis), identified using Hotspot. Integrated modules are ordered by decreasing overlap and reveal unique combinatorial patterns.
**(f)** Spatial distribution of representative integrated modules (e.g., Module 14 and Module 17), highlighting spatially restricted expression patterns.
**(g)** Pathway enrichment analysis of modules from h. Module 14 is enriched for immune activation and inflammatory pathways, while Module 17 is associated with interferon signaling and apoptosis, reflecting niche-specific functional states.
All scale bars: 500 µm.

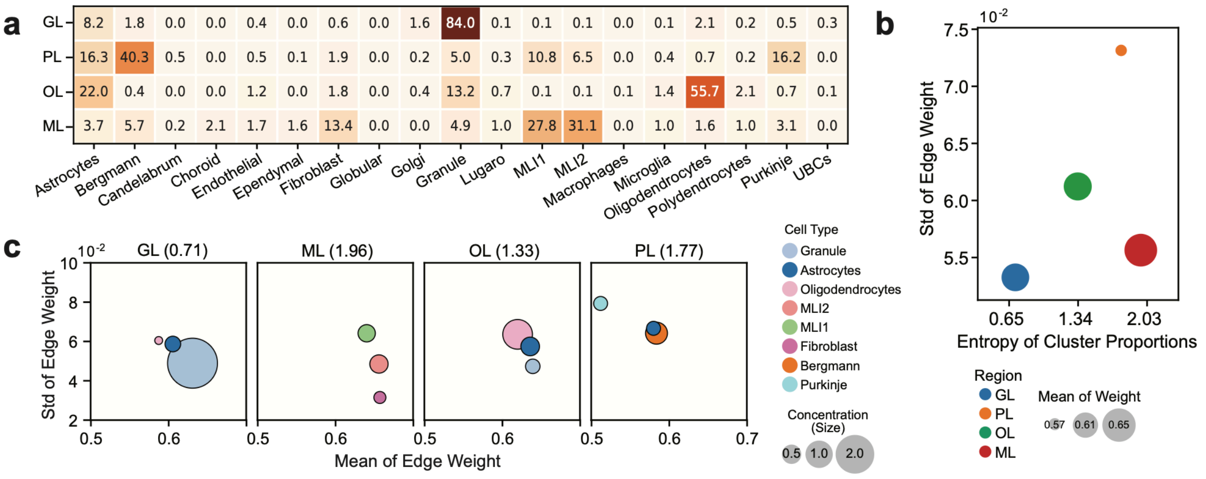


**Supplementary Figure 3 | Glimmer captures biologically structured neighbor graphs aligned with cerebellar architecture.**
**(a)** Cell type compositions across Glimmer-identified anatomical regions in Slide-seq cerebellum, including the granule layer (GL), Purkinje–Bergmann layer (PL), oligodendrocyte layer (OL), and molecular layer (ML). Values indicate the proportion (%) of each cell type within each region.
**(b)** Scatter plot showing the relationship between cell type entropy (x-axis) and the standard deviation of neighbor edge weights (y-axis) across cerebellar regions. Dot size

represents the mean edge weight within each region, and color indicates region identity as in panel (a).

**(c)** Cell-type-specific neighbor connectivity properties across regions. Each panel corresponds to one region (entropy shown in parentheses). Dots represent the top-3 dominant cell types per region. The x-axis shows the mean edge weight, and the y-axis shows the standard deviation (scaled ×100). Dot color indicates cell type; size reflects relative concentration (abundance normalized by regional entropy).

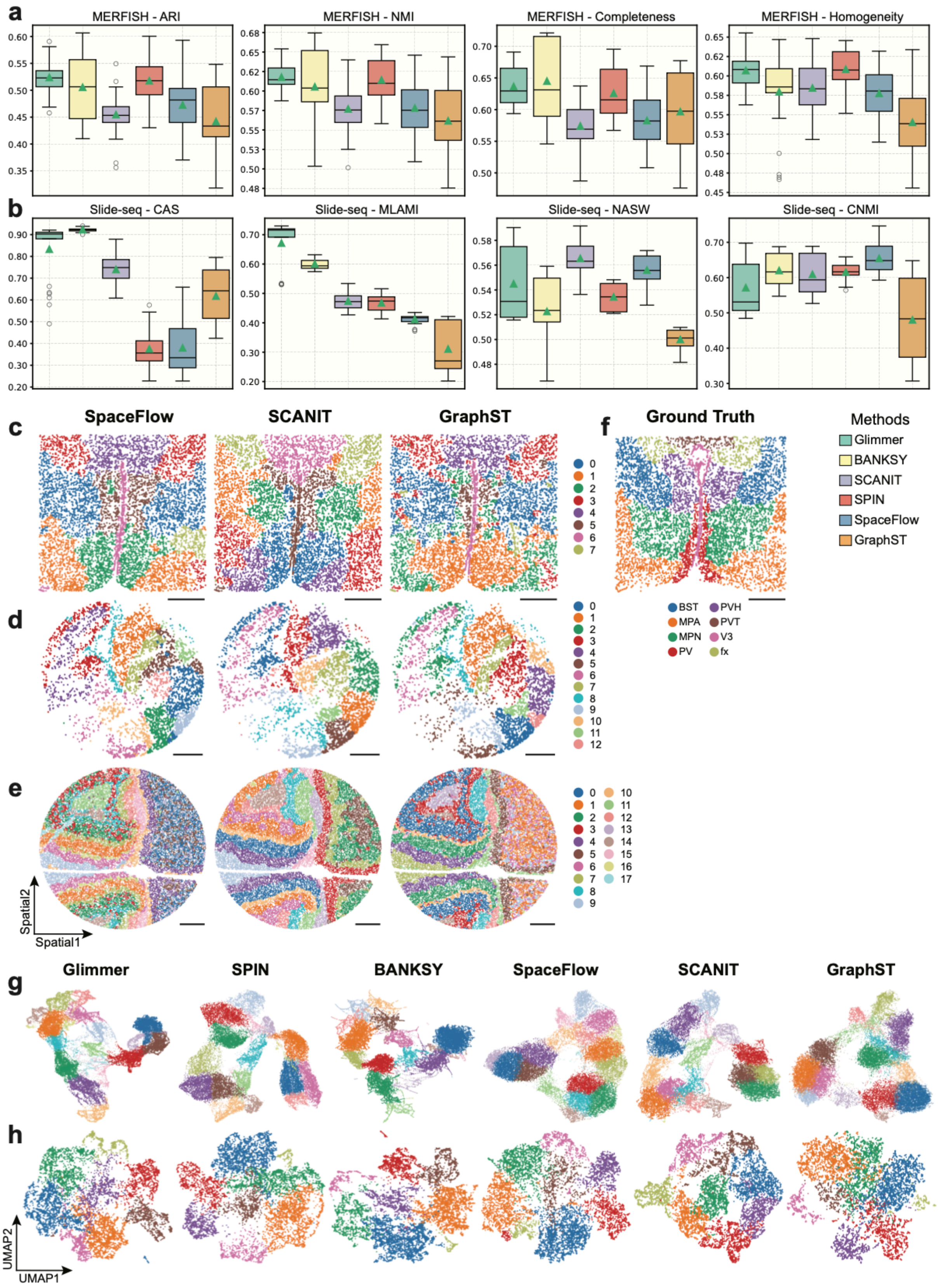


**Supplementary Figure 4 | Glimmer generalizes spatial domain identification**

**across scales and technologies.**

**(a-b)** Benchmarking of six spatial domain identification methods (Glimmer, SPIN, BANKSY, SpaceFlow, SCANIT, GraphST) on five region-annotated MERFISH datasets (a) and five unlabeled Slide-seq datasets (b), using supervised metrics (ARI, NMI, Completeness, Homogeneity) and unsupervised metrics (CAS, MLAMI, NASW, CNMI), respectively. Each method was run five times with different random seeds.

**(c-e)** Predicted spatial domains across representative methods for MERFISH hippocampus (c), Slide-tags mouse brain (d), and Slide-seq cerebellum (e), showing method-specific domain assignments across platforms.

**(f)** Ground truth for MERFISH hippocampus domain segmentation.

**(g-h)** UMAP of predicted spatial domains from all methods for Slide-seq cerebellum (g) and MERFISH hippocampus (h).

All scale bars: 500 µm.

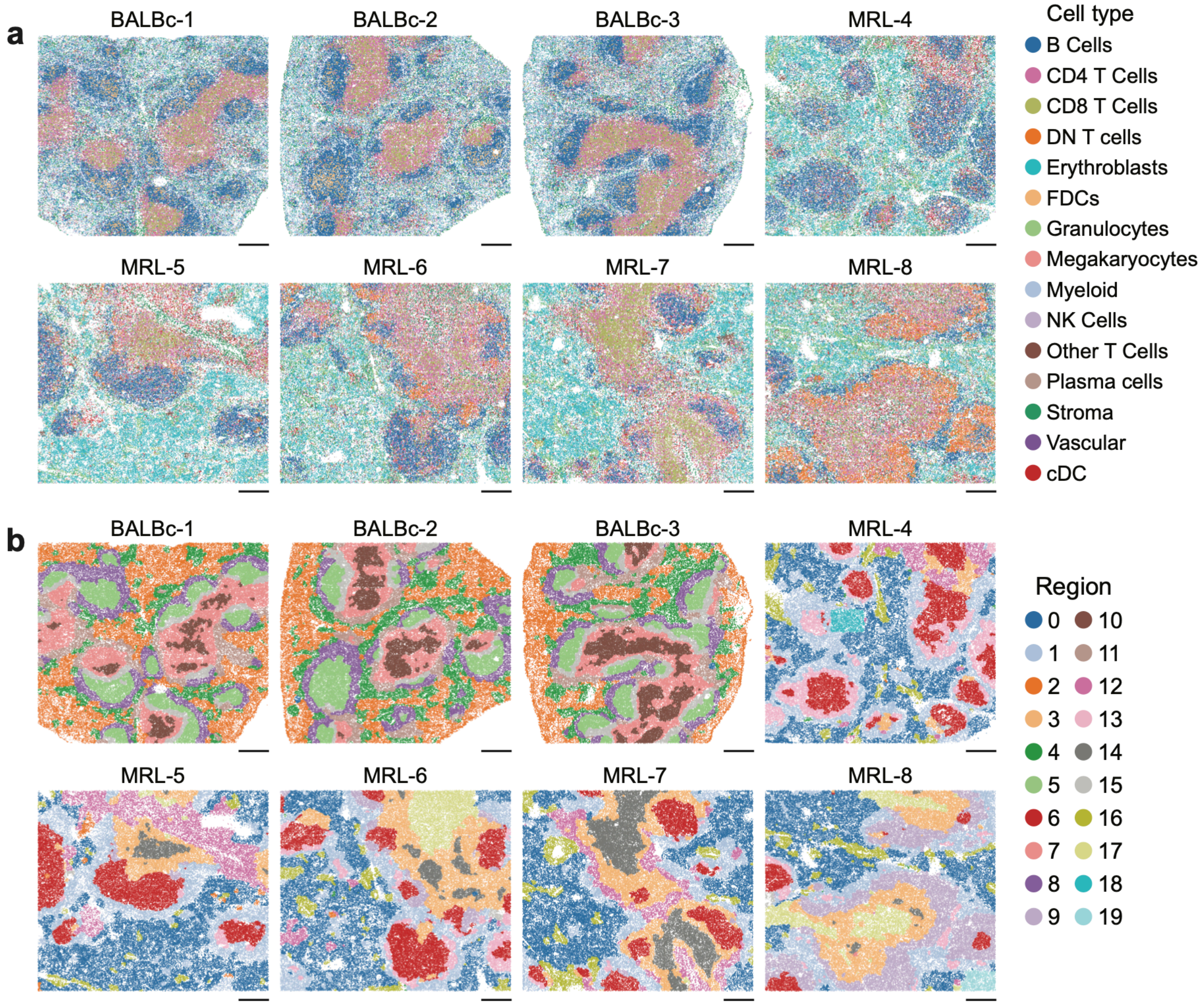


**Supplementary Figure 5 | Glimmer enables consistent identification of protein-level spatial domains across CODEX spleen sections.**

**(a)** Cell type annotations across spleen sections from three healthy BALB/c and six lupus-prone MRL/lpr mice.

**(b)** Glimmer-derived integrative spatial domains reveal conserved architectures in health and altered organization in disease.

All scale bars: 250 μm.

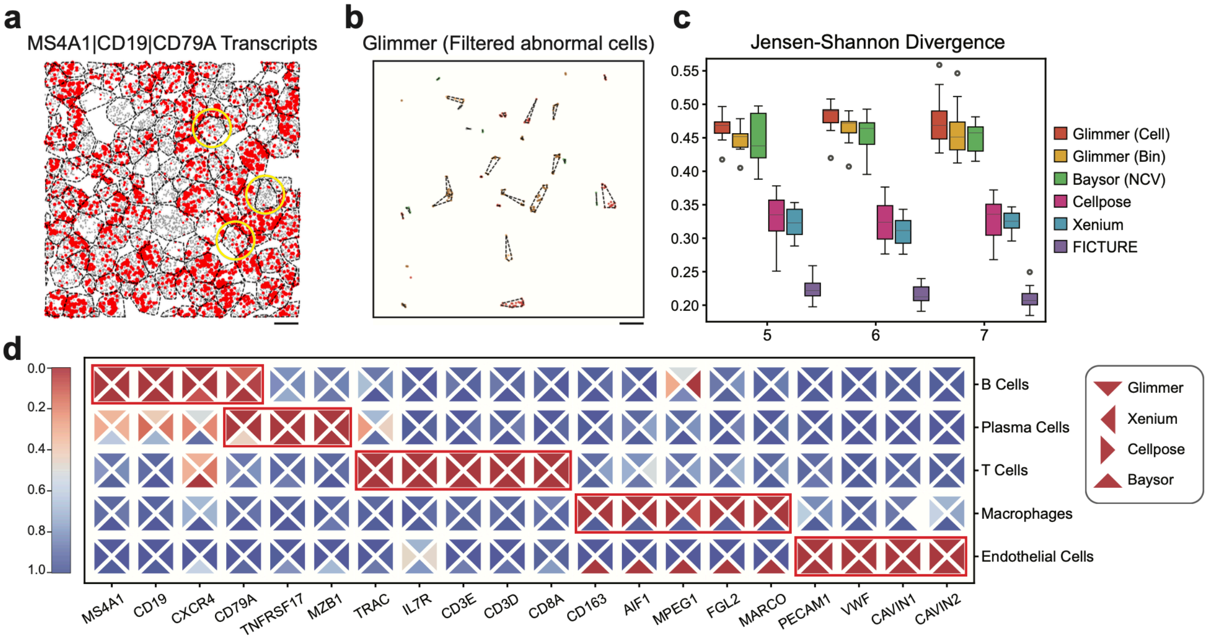


**Supplementary Figure 6 | Glimmer enables accurate transcript assignment and improves cluster purity.**

**(a)** Spatial distribution of B cell marker transcripts (*MS4A1*, *CD19*, *CD79A*) overlaid with Xenium-based cell boundaries. Dot size reflects local transcript density smoothed over a 15-neighbor kernel.

**(b)** Abnormal cell fragments filtered by Glimmer following Voronoi-based segmentation and transcript clustering refinement.

**(c)** Jensen-Shannon divergence of cluster-averaged gene expression across 15 Xenium patches (800 μm × 1600 μm), evaluated at cluster numbers of 5, 6, and 7, with five random seeds per setting. Higher values indicate greater inter-cluster separability. Box plots display the median (center line), interquartile range (box), and 1.5× IQR (whiskers), with outliers shown as individual points.

**(d)** Expression patterns of key marker genes across major immune and stromal cell types. Each triangle represents gene expression detected by a specific method (Glimmer, Xenium, Cellpose, or Baysor), with triangle orientation indicating the method. Expression values were scaled to the range [0, 1] within each method.

All scale bars: 2 μm.

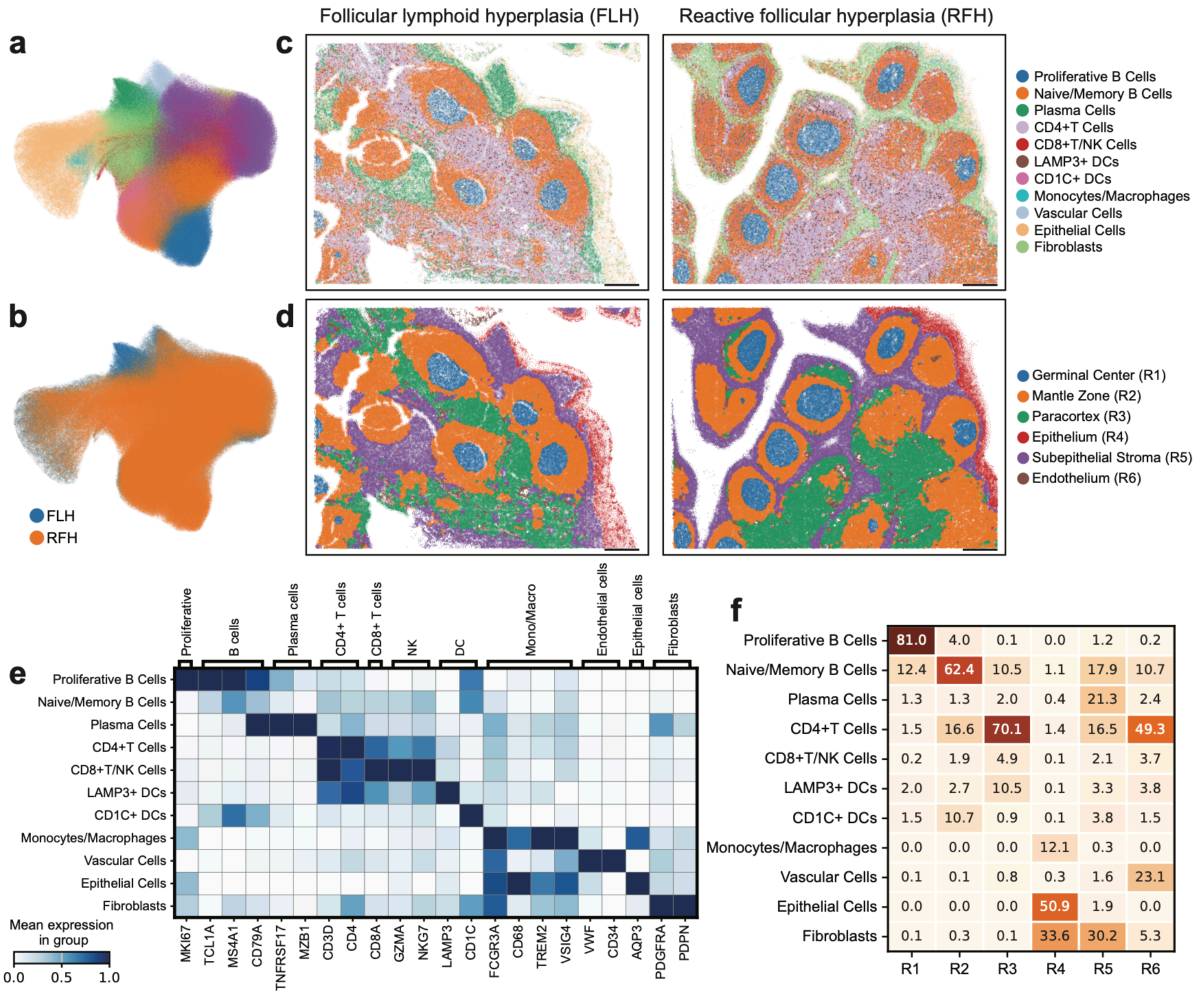


**Supplementary Figure 7 | Glimmer enables integrative clustering of cells and regions across pathological conditions.**

**(a)** UMAP visualization of cell types across integrated FLH (Follicular Lymphoid Hyperplasia) and RFH (Reactive Follicular Hyperplasia) samples, with cell type labels consistent with panel (g).

**(f)** Batch-corrected UMAP embedding after scVI integration, colored by condition.

**(g)** Spatial mapping of predicted cell types across tissue sections from FLH and RFH.

**(h)** Spatial segmentation of tissue regions (e.g., Germinal Center, Mantle Zone, Epithelium) based on transcript-level clustering.

**(i)** Mean expression heatmap of marker genes across annotated cell types shown in panels (e) and (g).

**(j)** Cell type compositions across spatial regions defined in panel h, highlighting enrichment patterns such as proliferative B cells in germinal centers (R1) and fibroblasts in subepithelial stroma (R5).

All scale bars: 250 µm.
